# Widespread attraction of marine bacteria to the macroalgae-derived polysaccharide fucoidan

**DOI:** 10.64898/2026.10.08.757582

**Authors:** Valerio Anelli, Jean-Baptiste Raina, Riccardo Foffi, Linda Biehler, Richard J. Henshaw, Blake Ushijima, Margot Bligh, Giovanni Stefano Ugolini, Sam Charlton, Milad Radiom, Natasha Blitvic, Raffaele Mezzenga, Andreas Sichert, Jan-Hendrik Hehemann, Johannes Keegstra, Roman Stocker

## Abstract

Fucoidan is the second most abundant polysaccharide in brown macroalgae, contributing up to 50% of the dissolved organic matter released daily from these algae into the ocean, substantially influencing the marine carbon cycle. Due to its complex chemical structure, fucoidan is highly resistant to microbial degradation and accumulates on the surface of macroalgae, where it contributes to regulating their microbiome. Here, we used a range of microfluidic assays to quantify the chemotactic and physiological responses of marine bacteria to fucoidan. Our results reveal that all the strains tested (14 out of 14) are attracted to fucoidan produced by two different species of brown macroalgae. Attraction occurred only at high fucoidan concentrations (0.1 to 1 mg/mL), such as those expected in the immediate vicinity of algal surfaces, whereas low concentrations elicited no response or modest chemorepulsion. Although fucoidan did not support bacterial growth under our experimental conditions, it did not impair the utilization of other carbon sources. Together, these findings suggest that fucoidan gradients can act as chemical cues guiding marine bacteria toward the nutrient-rich microenvironment of macroalgal surfaces, pointing to a potential ecological role of fucoidan in shaping macroalgal microbiota, bacterial behavior, and ultimately carbon cycling in the ocean.

## Introduction

Macroalgal glycans represent a major source of organic carbon in marine environments^1,2^, accounting for up to a third of the dissolved (DOC) and particulate (POC) organic carbon pools in macroalgae-dominated systems ^3^. Consequently, they play a central role in ocean carbon cycling and sequestration^4,5^. Fucoidan is one of the predominant polysaccharides in brown macroalgae^6^, representing up to 30% of their dry weight^7^. It serves key structural and protective functions on the walls of algal cells^8^ and constitutes up to 50% of the dissolved organic matter exuded daily by brown macroalgae^9^, resulting in the release of a large amount of carbon into the ocean^10^. Given its recalcitrant nature, fucoidan is likely to accumulate as both high-molecular-weight dissolved organic matter (DOM) and particulate organic matter (POM) in the ocean water column. This accumulation contributes to long-term carbon sequestration by fueling the biological carbon pump through export to the deep sea and burial in coastal sediments^11,12,9^.

The biochemical composition of the surface of brown macroalgae^9^ has been proposed to affect the composition of host-associated microbial communities^13,14^. Chemical interactions within the macroalgae diffusive boundary layer play a central role in shaping the macroalgae holobiont^15^, as both bacteria and their hosts rely on the reciprocal exchange of metabolites that support growth, defense, and nutrient acquisition^16^. A wide range of low–molecular-weight compounds, including amino acids, sugars, and secondary metabolites, regulate these interactions by attracting, repelling, or nourishing associated microbes^17^. However, the mechanisms through which large polysaccharides such as fucoidan contribute to microbial assembly and influence microbial behavior at the macroalgal surface remain largely unexplored.

Fucoidans comprise a diverse family of heavily branched, sulfated bioactive polysaccharides with complex chemical structures, primarily built around a fucose backbone^18^. As high–molecular-weight recalcitrant polymers^19^, fucoidans exhibit low diffusivity and slow turnover^20^, physicochemical properties that are expected to result in strong, persistent chemical gradients in the immediate vicinity of algal surfaces. The structure of fucoidan varies widely among producing species and environmental conditions, resulting in substantial differences in sulfate content, molecular weight, and monosaccharide composition ^21,19,22^. This biochemical heterogeneity and complexity underpin fucoidan’s notable resistance to microbial degradation: only a limited number of highly specialized, particle-associated bacterial taxa can partially degrade these polymers using dedicated carbohydrate-active enzymes ^23,24,25^. These same biochemical features also give origin to a wide range of bioactive functions^26,27,28,29^, including antimicrobial activity^30,31,32,33^, which have been reported predominantly in clinical bacteria under laboratory conditions^32,34^.

In the ocean, bacterial chemotaxis can regulate access to hosts and thereby control the assembly of marine microbiomes^35,36,37^. A broad range of molecules can act as signaling cues, though most of them are small molecules and polysaccharides have been largely overlooked^38^. In this context, recent work has revealed that different glycans, including laminarin^39^, alginate^39,40^, and pectin^40^, can act as potent chemoattractants in the ocean. These findings suggest that chemotaxis toward polysaccharides may be a widespread microbial behavior in natural environments^41,42^. In contrast, the pronounced structural recalcitrance of fucoidan^24^ and its reported antibacterial activity^32^ would go against a potential role as a chemical cue for marine bacteria. Here, we investigate how fucoidan influences the behavior of marine bacteria. Using a range of microfluidic behavioral assays^43,44,45^, we quantify the chemotactic response to fucoidan across a range of environmentally relevant concentrations for 14 marine bacteria spanning different taxa, habitats, lifestyles, and typical exposures to algal-derived organic matter, and measure the effect of fucoidan on their growth. We find that high concentrations of fucoidan reliably elicit strong chemoattraction across all of marine bacteria, for at least one of the fucoidan obtained from two different algal species, pointing to a new signaling role of fucoidan in the ocean. Contrary to expectations, fucoidan did not exhibit any detrimental effects on the marine strains tested when additional carbon sources were present, indicating that many bacteria can also grow in fucoidan-rich environments. Together, these findings identify a previously unrecognized ecological role for fucoidan as a chemical cue that structures macroalgae–bacteria interactions through chemotaxis, with implications for microbial foraging behavior and the cycling of algae-derived carbon in marine ecosystems.

## Results

### Fucoidan elicits a strong chemotactic response in marine bacteria

To investigate the chemotactic response of marine bacteria to fucoidan, we used the In Situ Chemotaxis Assay (ISCA)^43,44,39,35^, which is composed of an array of 20 microwells that can be filled with different chemoeffectors. Once the device is deployed in an aqueous environment, the chemoeffector from each well diffuses out through a small inlet and forms a concentration gradient that attracts into the well bacteria that are chemotactic to that chemical. Bacteria in each well can subsequently be collected and enumerated by flow cytometry. Using the ISCA, we quantified the behavior of 14 motile marine strains belonging to three different categories: (i) 4 strains previously found to chemotax toward other polysaccharides (laminarin and alginate) ^39^, (ii) 3 strains from the genus *Vibrio* that have been well studied for their motility and chemotactic responses toward other molecules ^46,47^, and (iii) 7 strains isolated from brown-macroalgae forests on the North-west Atlantic coast^48^. Strain selection prioritized isolates with confirmed motility, as chemotaxis requires motility. Many known fucoidan-degrading strains were excluded from testing based on a genomic screen of a fucoidan-associated bacterial community (Fig. S1; Methods), which revealed that 7 out of 8 degraders lack a complete flagellar biosynthesis and chemotaxis gene set. We performed ISCA experiments with this strain collection using fucoidan obtained from two different algal species: *Fucus vesiculosus* and *Laminaria hyperborea*. The concentration range tested (0.001-1 mg/mL) was selected to be ecologically relevant by computing the fucoidan concentration expected on the algal surface from measurements of the fucoidan leakage rate (Supplementary Note 1), obtained by extracting and quantifying the amount of fucoidan accumulated on the surface of a *F. vesiculosus* specimen in a one-hour incubation in water. As a quantitative measure of the strength of chemotaxis, we used the chemotactic index^43^, *I_C_*, computed as the ratio of the number of cells in the treatment wells containing fucoidan (dissolved in filtered artificial seawater, ASW), to the average number of cells in the control wells containing fucoidan-free ASW. Thus, an *I_C_* value significantly larger than 1 denotes chemoattraction, whereas a value significantly lower than 1 denotes chemorepulsion.

We first tested chemotaxis for the 14 bacterial strains to commercially available fucoidan from *F. vesiculosus* (Sigma; See Methods). We found that 11 of the 14 strains exhibited chemoattraction to the highest concentration of fucoidan (1 mg/mL) from *F. vesiculosus*, with *I_C_* values ranging from 2 to 140 (Fig. 1A, Dunnett’s test, *p <* 0.05, Supplementary Data 1): across all strains tested, chemoattraction to this concentration was highly significant (t-test, *p* = 0.0005, Supplementary Data 1). *Vibrio coralliilyticus* sp. YB1, *Pseudoalteromonas* sp. ASV109 and *Shewanella* sp.AS82 were also attracted to 0.1 mg/mL fucoidan from *F. vesiculosus*, while no strain was attracted to concentrations lower than 0.1 mg/mL of this fucoidan. Instead, three strains showed chemorepulsion at low fucoidan concentrations, particularly at 0.001 mg/mL, with *I_C_* values ranging from 0.3 to 0.6 (Fig. 1A, Dunnett’s test, *p <* 0.05, Supplementary Data 1): across all strains tested, chemorepulsion from this concentration was significant (t-test, *p* = 0.01, Supplementary Data 1). Notably, *Pseudoalteromonas* sp. AS80, *Pseudoalteromonas* sp. AS81 and *Pseudoalteromonas* sp. ASV16 exhibited a biphasic chemotactic response, showing attraction at high concentrations and repulsion at low concentrations of fucoidan from *F. vesiculosus*.

**Figure 1:**
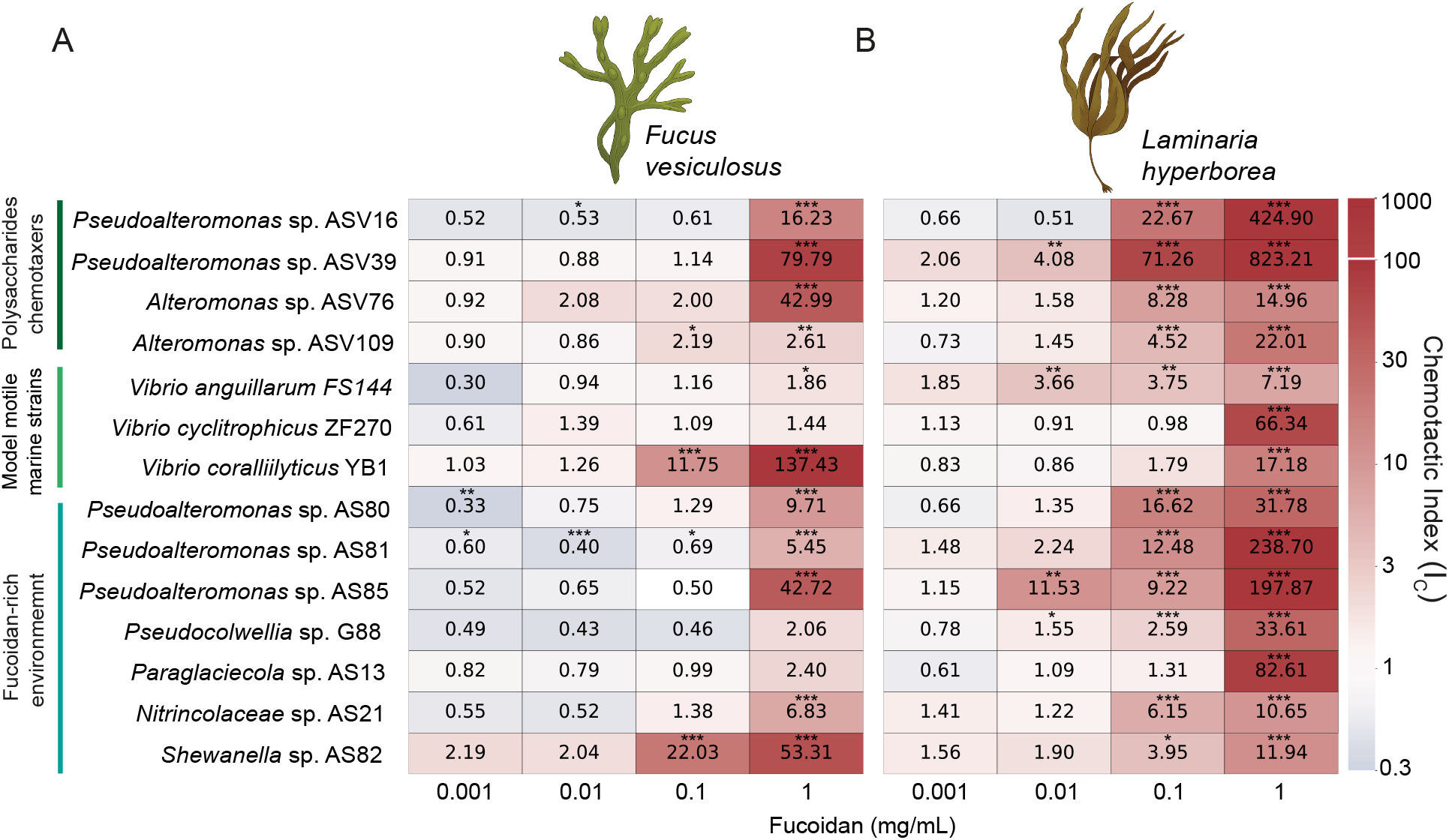
Bacterial chemotactic response to fucoidan from *F. vesiculosus* and *L. hyperborea*. **(A-B)** Heatmap summarizing the chemotactic responses of 14 bacterial strains (rows) toward different concentrations of fucoidan (columns) from two macroalgal species. Values represent the average chemotactic index (*I_C_*) relative to the artificial seawater (ASW) control (*n* = 3). Stars indicate chemotactic responses significantly higher (attraction) or lower (repulsion) than to the control ASW (* = *p <* 0.05; ** = *p <* 0.01; *** = *p <* 0.001; Dunnett’s test, with Benjamini–Hochberg p-value adjustment for testing across multiple strains). All *p* values are reported in Supplementary Data 1.

Fucoidan is highly polydisperse^49^, being composed of a mixture of chains of different lengths and molecular weights^30,32^. To ensure that the observed chemotactic responses were not due to small-molecular-weight contaminants, we first purified the crude fucoidan source via anionic exchange chromatography (AEX; Methods) to remove non-charged contaminants, such as neutral oligosaccharides. This purified fucoidan was tested on a subset of six strains from the three different groups (Fig. S2). While the purification process caused a general reduction in *I_C_* values (from an average of 30 to 10) at high fucoidan concentration (1 mg/mL), the chemotactic responses were consistent with those observed prior to purification, with all six strains showing significant attraction at 1 mg/mL (Fig. S2, Dunnett’s test, *p <* 0.05, Supplementary Data 2). In contrast, after purification, significant chemorepulsion at 0.001 mg/mL was present only in *Pseudocolwellia* sp. G88 (Fig. S2, Dunnett’s test, *p <* 0.05, Supplementary Data 2), showing that low-concentration repulsion was not a general feature of the response to fucoidan. Because AEX separates by charge rather than by size, it cannot alone exclude the presence of small contaminants. We therefore performed size-fractionation of both AEX-purified and commercially available fucoidan using Amicon centrifugal filters (5 kDa molecular-weight cutoff; Methods), and tested 1 mg/mL of the resulting high- and low-molecular-weight fractions for chemotactic activity in four strains (Fig. S3). The high-molecular-weight fraction retained strong chemoattraction for both fucoidan sources (*I_C_* = 4.2–12.2, Dunnett’s test, *p <* 0.001, Supplementary Data 2), closely matching the response of the corresponding unfractionated preparations, whereas the low-molecular-weight fraction showed no response in three of four strains and only a modest response in the fourth (*Pseudoalteromonas* sp. AS85; *I_C_* = 2.3; Dunnett’s test, *p <* 0.05, Supplementary Data 2). Additionally, the washout controls (ASW used to wash the retained high-molecular-weight fraction during fractionation) showed no response across any strain or condition, ruling out any contribution of fractionation artifacts to the chemotactic response. We further verified the structural composition of fucoidan across all preparations used in our assays by ^1^H-NMR (Fig. S4; Methods), confirming that the expected fucoidan structural signature (anomeric, ring, O-acetyl, and methyl proton resonances) dominates each preparation. Minor, unassigned peaks were present at low intensity in each preparation. Still, their pattern differed across sources, with no single peak shared by all four, thus excluding the presence of a common contaminant across preparations. These results indicate that the chemoattraction to high fucoidan concentrations is not caused by small contaminants potentially present in the different preparations tested, but rather by the fucoidan molecule itself.

To confirm the generality of chemoattraction toward high fucoidan concentrations, we tested the 14 strains for chemotaxis to fucoidan extracted from another well-characterized, widespread brown alga species, *L. hyperborea* ^50^, purified using AEX (Fig. 1B). In line with the response to fucoidan from *F. vesiculosus*, all strains exhibited chemoattraction to the highest concentration of fucoidan (1 mg/mL; Fig. 1B, Dunnett’s test, *p <* 0.05, Supplementary Data 1), with *I_C_* values that were even higher and up to 800 (more than 7-fold higher than any other chemoattraction ever recorded with the ISCA). Additionally, 11 of 14 strains were attracted to an intermediate concentration of fucoidan from *L. hyperborea* (0.1 mg/mL), with *I_C_* values between 2.5 and 70 (Fig. 1B, Dunnett’s test, *p <* 0.05, Supplementary Data 1): across all 14 strains, chemoattraction toward this concentration was significant (t-test, *p* = 0.00024, Supplementary Data 1). No strain exhibited chemorepulsion to any concentration of *L. hyperborea* fucoidan (Fig. 1B, Dunnett’s test, Supplementary Data 1). These results reveal that attraction to fucoidan from different macroalgae is prevalent among marine bacteria, is often strong or very strong, with some variability in the magnitude of the attraction associated with the specific fucoidan source. Moreover, the response was consistently of chemoattraction across all three groups, indicating a broadly conserved behavior among the strains tested, whereas chemorepulsion was substantially weaker and less robust and appeared to depend on the fucoidan source and purification procedure.

### Attraction toward fucoidan is mediated by the chemotaxis signaling pathway

To further confirm that the accumulation of cells in the ISCA wells was due to chemotaxis, we sought to exclude the possibility that the accumulation resulted from the physical properties of fucoidan. The first possibility we considered was a difference in viscosity between fucoidan solutions and seawater, which could have resulted in cells slowing down in the entrance of the wells due to the higher viscosity, and thus accumulating there even in the absence of chemotaxis^51^. To test this, we measured the viscosity of different concentrations of commercially available fucoidan from *F. vesiculosus*, from 1 to 10 mg/mL, via rheometry (Fig. S5; Methods): we found no significant differences in viscosity between the control seawater and the fucoidan solutions (Fig. S5; Two-sample t-test, Supplementary Data 2). Furthermore, we quantified the swimming speed of *Pseudoalteromonas* sp. ASV16, a species that exhibited high chemotactic indices in the ISCA experiments (Fig. 1, Fig. S2; Fig. S3), by tracking cells in the presence of uniform concentrations of commercially available fucoidan from *F. vesiculosus*, ranging from 0.001 to 10 mg/mL. These experiments showed that the presence of fucoidan did not reduce either the fraction of swimming cells or the cells’ swimming speed (Fig. S6), further ruling out the physicochemical properties of the fucoidan solution as being potentially responsible for trapping cells in the ISCA wells. Rather, cell tracking showed that the presence of fucoidan caused an increase in the average swimming speed from 35 to 50 *µ*m s*^−^*^1^ (Fig. S6), indicating a positive chemokinetic response to fucoidan.

We performed additional experiments to determine whether the accumulation of bacteria in ISCA wells was a genuine chemotaxis response mediated by the chemotaxis signaling pathway. Many of the marine isolates used in this study are currently not genetically tractable, except *Vibrio coralliilyticus* YB1, for which we constructed a non-chemotactic mutant by deleting the signaling kinase (ΔCheA)^52^ (see Methods). We then exposed both the wild-type (WT) and the non-chemotactic (ΔCheA) mutant to AEX-purified fucoidan from *F. vesiculosus* (used in Fig. S2), at multiple concentrations, in parallel ISCA experiments performed on the same day (Fig. 2A). The response of the non-chemotactic mutant to all fucoidan concentrations tested was indistinguishable from its response to the ASW control, with *I_C_* values very close to 1 (from 0.9 to 1.3; Fig. 2A, Dunnett’s test, *p <* 0.05, Supplementary Data 1). In contrast, WT *Vibrio coralliilyticus* YB1 displayed *I_C_* values ranging from 4 to 13 in these experiments (Fig. 2A, Dunnett’s test, *p <* 0.05, Supplementary Data 1). These results confirm that the accumulation of cells in ISCA wells is mediated by the chemotaxis pathway and thus is a genuine chemotactic response.

**Figure 2:**
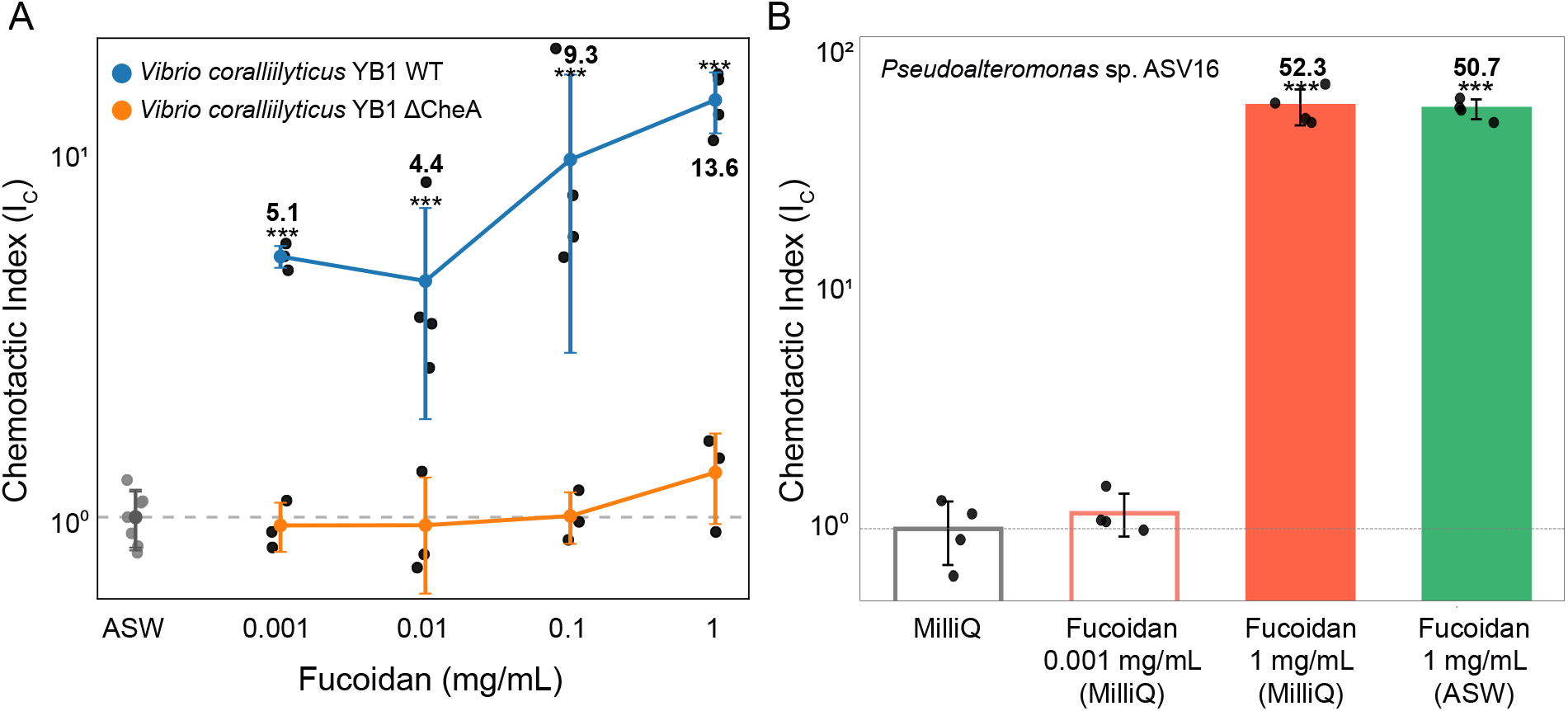
Bacterial accumulation in wells containing fucoidan is mediated by the canonical chemotaxis pathway. **(A)** The chemotactic index of the non-chemotactic mutant (ΔCheA) and the wild type (WT) *Vibrio coralliilyticus* YB1 to different concentrations of AEX-purified fucoidan from *F. vesiculosus*. The horizontal dashed gray line indicates the control *I_C_* = 1 value. Error bars indicate standard deviations across replicates (*n* = 3) and stars indicate significant chemoattraction compared to the control (ASW; Dunnett’s test, *** = *p <* 0.001) with corresponding average significant *I_C_* values reported on top of the lineplot. **(B)** Chemotactic response of *Pseudoalteromonas* sp. ASV16 to 1 mg/mL of AEX-purified fucoidan in Milli-Q water supplemented with NaCl (35 g/L) and in artificial seawater (ASW). The horizontal dashed gray line indicates the control *I_C_* = 1 value of supplemented Milli-Q. Stars and filled barplots indicate significant chemoattraction compared to the control (Dunnett’s test, *** = *p <* 0.001) with corresponding average significant *I_C_* values reported on top of the barplots. All *p* values are reported in Supplementary Data 1.

We lastly investigated the possibility that the attraction to fucoidan is mediated by counterion gradients that are associated with fucoidan dissolved in seawater. Seawater is a highly ionic solution and ions are ubiquitous in the ocean and typically present at millimolar concentrations^53^; therefore, any charged species, including fucoidan, is accompanied by a diffuse layer of associated counter-ions^54^. Such counter-ions could influence bacterial chemotactic responses and potentially mediate attraction if they themselves act as chemoeffectors. Several inorganic ions elicit repellent responses in bacteria^55,56^ and, for metals, chemoattraction has been reported primarily toward reduced forms rather than the oxidized forms that dominate in marine environments^57^. More recently, chemotaxis toward potassium ions has been reported in *Escherichia coli* ^58^. To test for a potential contribution of counter-ions to the observed chemoattraction toward fucoidan, we measured the chemotactic response of *Pseudoalteromonas* sp. ASV16, one of the strains that exhibited the strongest attraction to 1 mg/mL fucoidan, using polymer solutions with and without associated counter-ions (Fig. 2B). AEX-purified fucoidan from *F. vesiculosus* was dissolved in ion-free Milli-Q water or in ion-rich artificial seawater (ASW) as a control. To permit motility in ion-free conditions, Milli-Q water was supplemented with seawater concentrations of sodium (NaCl, 35 g/L), as flagellar motility in many marine bacteria is powered by a sodium-motive force^59^. In both ion-free and ion-rich conditions, *Pseudoalteromonas* sp. ASV16 exhibited a strong attraction to fucoidan (Fig. 2B; Dunnett’s test, *p <* 0.05; Supplementary Data 1), ruling out the possibility that the observed chemotactic responses were due to counter-ions.

Together, these results demonstrate that fucoidan induces a behavioral response in motile marine bacteria. This behavior is caused by the polymer itself and mediated by the chemotaxis pathway, and most marine bacteria tested are highly attracted to high fucoidan concentrations from multiple macroalgal sources.

### Chemotaxis to fucoidan occurs in response to steep gradients

The ISCA is ideally suited to investigate the chemotactic response of bacteria to multiple compounds or concentrations, but it does not capture the temporal dynamics of the response, rather providing a single endpoint measurement. To investigate the dynamics of chemotaxis to fucoidan, we complemented the ISCA results with experiments using a second microfluidic assay. We used an established three-inlet gradient generator ^45,60^ that allowed us to visualize the temporal evolution of the bacterial chemotactic response to a fucoidan pulse with high temporal resolution (1 frame per second) over a 10-min period (Methods). We injected a fucoidan solution, a cell suspension, and ASW through the three inlets of this gradient generator device to form three adjacent flowing streams having a 4:1:4 thickness ratio (Fig. 3A). Upon stopping the flow and imaging the dynamics at a fixed location in the microchannel, fucoidan diffuses from its original stream into the two adjacent streams, establishing a gradient that weakens over time^61,62,45^. During this process, time-lapse imaging is applied, and the positions of individual bacteria are then extracted using standard image analysis and particle location techniques (Methods). It is important to note that the timescale over which the gradient evolves in the gradient generator device is much shorter than in the ISCA, where the reservoir (the fucoidan originally in the well) is much larger, thus sustaining a gradient for much longer.

**Figure 3:**
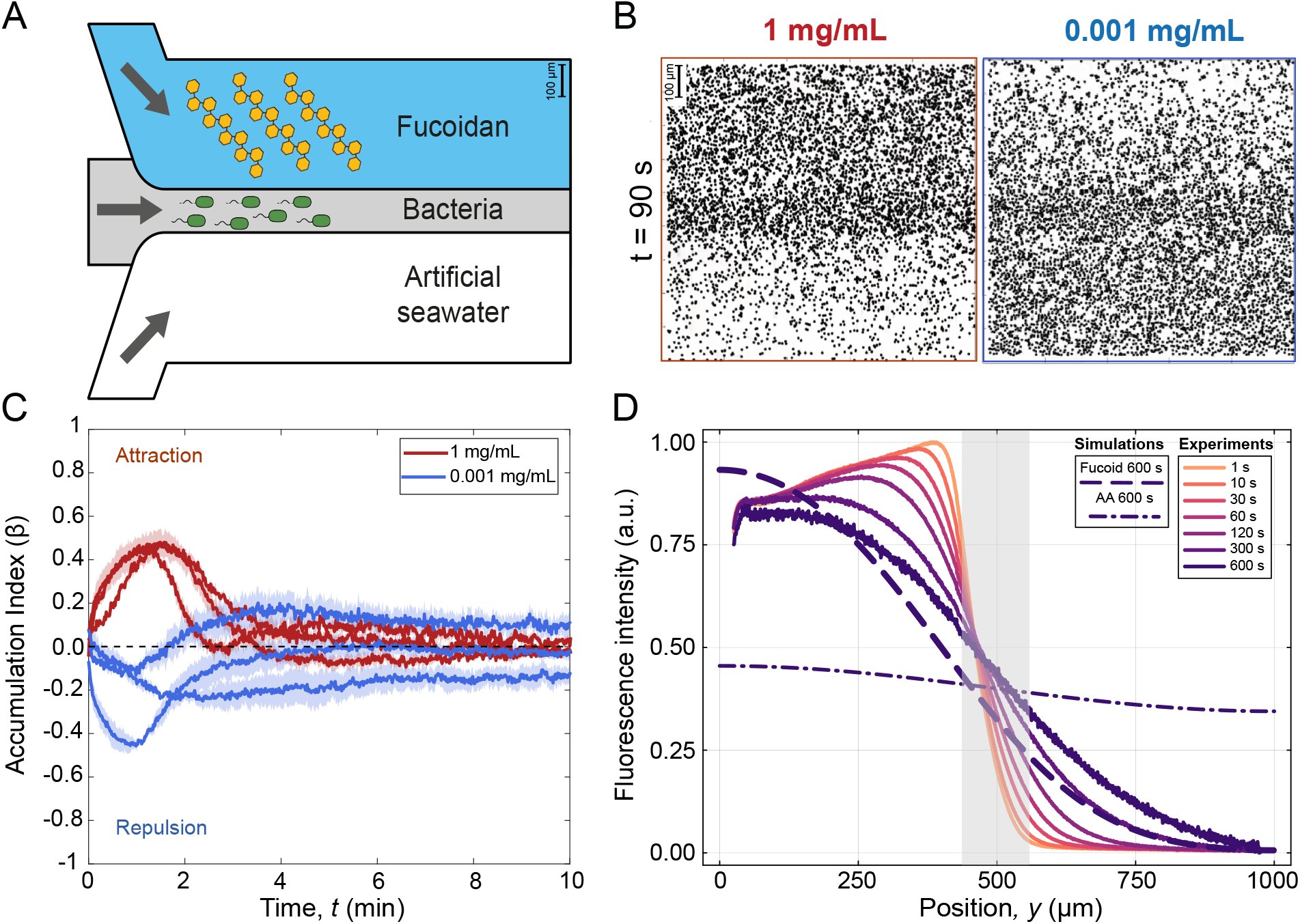
Microfluidic characterization of the chemotactic response of *Pseudoalteromonas* sp. ASV16 to fucoidan gradients. **(A)** Schematic of the microfluidic gradient generator, with the chemoeffector (AEX-purified fucoidan from *F. vesiculosus* suspended in ASW) on one side of the channel, bacteria suspended in ASW in the center, and ASW on the other side. **(B)** Snapshot of bacterial positions across the width of the channel for high (red, left) and low (blue, right) concentrations of fucoidan 90 s after stopping the flow, corresponding to the peak chemotactic response (see panel C). Each black dot represents a single bacterium. Videos of the time course of the chemotactic response are included in the Supplementary Material (Supplementary Videos 1 and 2). **(C)** Quantification of the chemotactic response in terms of the β accumulation index in response to high (red) and low (blue) concentrations of fucoidan as a function of time. β *>* 0 and β *<* 0 denote attraction and repulsion, respectively. Each line represents a biological replicate and comprises three technical replicates; the shaded region shows the standard deviation among the technical replicates (n=3). **(D)** Fucoidan concentration profiles across the width of the microfluidic channel measured using fluorescently labelled fucoidan (colors indicate different time points, see legend), overlaid with simulated profiles of fucoidan (dashed line) and an amino acid (dash-dotted line) at the final time of *t* = 600 s. Simulations were performed using approximated diffusivities of 40 *µ*m^2^ s*^−^*^1^ for fucoidan and 400 *µ*m^2^ s*^−^*^1^ for the amino acid. Concentration values are reported normalized by the initial concentration, *C*_0_. Fluorescence intensity was normalized to the initial concentration of fucoidan. The vertical gray band denotes the position of bacteria at time zero.

We used this gradient generator to measure the chemotactic response of *Pseudoalteromonas* sp. ASV16 to different concentrations of AEX-purified fucoidan from *F. vesiculosus* and *L. hyperborea*. The response was quantified using an accumulation index, β(*t*), defined as the difference in the number of cells within 200 *µ*m of the upper and lower bounds of the microchannel, normalized by the total cell count in these two regions at time *t* (Methods). A positive/negative value of β(*t*) indicates accumulation/repulsion, respectively. When we exposed *Pseudoalteromonas* sp. ASV16 to 1 mg/mL AEX-purified fucoidan from *F. vesiculosus* we observed a clear accumulation of cells toward the fucoidan region of the channel, with approximately 2.3*×* more cells in the upper region than in the lower one (β(*t*) = 0.4; Fig. 3B left; Fig. 3C red; Supplementary Video 1), consistent with the ISCA results (Fig. S2). At low fucoidan concentrations (0.001 mg/mL), we observed a chemorepulsion away from fucoidan (β(*t*) = *−*0.2; Fig. 3B right, Fig. 3C blue; Supplementary Video 2), in line with the low *I_C_* values (*≈* 0.5) measured with the ISCA. A similar pattern was observed with *Pseudoalteromonas* sp. ASV16 exposed to fucoidan from *L. hyperborea* at three concentrations: 1, 0.1 and 0.001 mg/mL. Consistent with the very high *I_C_* values exhibited by this strain when exposed to this fucoidan (Fig. 1), β(*t*) reached values close to the maximum of 1 (i.e., all motile cells had migrated to the upper 200 *µ*m of the channel; Fig. S7). At low fucoidan concentrations (0.001 mg/mL), *Pseudoalteromonas* sp. ASV16 cells were repelled (β(*t*) = -0.8), showing a clear biphasic response.

To better understand the dynamics of the bacterial response to fucoidan, we compared how the bacterial accumulation in the gradient generator related to the time-evolution of the gradient. Although the magnitude of the chemotactic response varied between the different concentrations tested, the response was consistently short-lived: cell accumulation peaked within 60 to 90 s after flow stoppage and relaxed to an almost uniform distribution across the channel width by 180 s (β(*t*) *≈* 0, Fig. 3B–C; Supplementary Videos 1 and 2). To directly quantify the time course of the fucoidan concentration profile across the channel, we imaged the evolution of the gradient from fluorescently labeled fucoidan from *F. vesiculosus* in the microfluidic channel (Fig. 3D). The observed gradient evolution matched the one predicted from a numerical simulation using the measured diffusion coefficient of fucoidan (Fig. S8, Fig. S9, Fig. 3D). These findings show that fucoidan gradients equilibrate much more slowly compared to those of small molecules, such as amino acids (Fig. 3D). Furthermore, they also confirm - as expected based on the low diffusivity of a molecule as large as fucoidan - that the short-lived behavioral response of bacteria to fucoidan cannot be explained by the disappearance of the fucoidan gradient due to diffusive equilibration. Instead, based on the 60 - 90 s response time of bacteria and considering the magnitude of the fucoidan gradient during this time window, we conclude that bacteria respond only to steep fucoidan gradients (0.01 *−* 0.005 mg*/*mL *µ*m*^−^*^1^; Fig. S10) and not to weak ones (*<* 0.005 mg*/*mL *µ*m*^−^*^1^; Fig. S10). This is consistent with the presence of strong gradients in the ISCA, as computed by numerical simulations of the ISCA (Fig. S11; Supplementary Note 2). Together, this analysis reveals that bacteria responding to fucoidan do so only when the fucoidan gradients are strong.

### High concentrations of fucoidan do not inhibit bacterial growth on different carbon sources

Our observation of widespread and strong chemotaxis toward fucoidan is surprising in view of fucoidan’s recalcitrance to bacterial consumption and its reported antibacterial effects^63,32,33^. To assess a potential bactericidal effect of fucoidan in our case, we tested the effect of commercially available fucoidan from *F. vesiculosus* on *Pseudoalteromonas* sp. ASV16 using propidium iodide (PI) staining to detect cell death (Methods). We used both microfluidic chambers, where we imaged cells exposed to high concentrations of fucoidan (1 mg/mL) for 48 h, and a bulk assay, in which we incubated cells with either high (1 mg/mL) or low (0.001 mg/mL) concentrations of fucoidan for 24 h and then quantified by flow cytometry (Fig. S12). Both assays showed no evidence of fucoidan-induced cell death (Fig. S12; two-sample t-test, *p <* 0.05; Supplementary Data 2). Moreover, we could not detect any signs of cell impairment, as bacteria retained vigorous motility throughout the entire microfluidic experiment (Supplementary Video 3). Next, we quantified the growth behavior of four strains from the three different ecological groups (*V. cyclitrophicus* ZF270, *Pseudoalteromonas* sp. ASV16, *Pseudocolwellia* sp. G88, and *Vibrio coralliilyticus* YB1) when exposed to commercially available fucoidan from *F. vesiculosus* as a sole carbon source. The bacteria were grown in a rich medium (Marine Broth), washed, and then exposed for 48 h to different concentrations of fucoidan (0.001, 0.01, 0.1, and 1 mg/mL) in ASW supplemented with f/2 nutrients, measuring the optical density (OD) every 10 minutes (Fig. 4A). When fucoidan was provided as the sole carbon source, the four strains exhibited only minimal growth, reaching OD_max_ of 0.02 after 48 h in 1 mg/mL of fucoidan (Fig. 4A, Fig. S13), only marginally higher than the initial inoculum (0.001) and in the same order as the background signal of the medium (0.005 to 0.01). This finding is in line with prior evidence that efficient fucoidan degradation is restricted to very few specialized strains, which belong to the *Verrucomicrobia* and *Planctomycetote* phyla ^23,24,64^.

**Figure 4:**
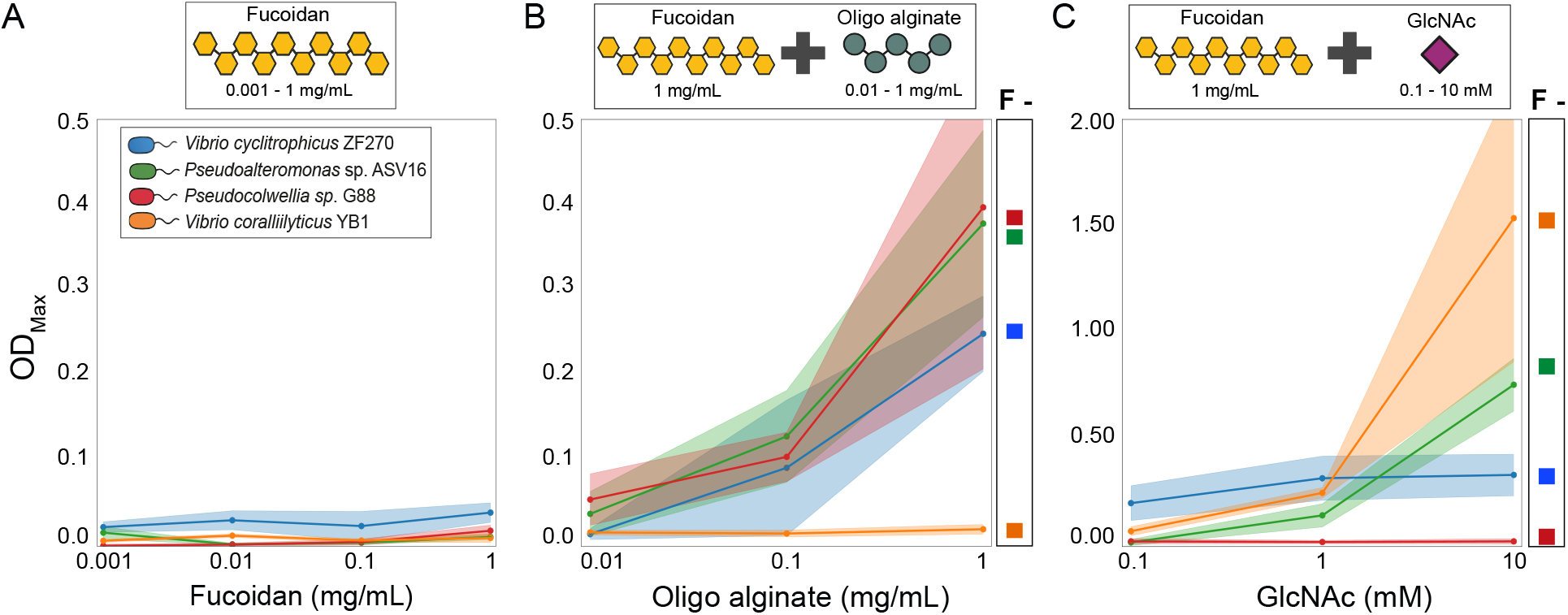
Growth of four marine strains on either fucoidan or two carbon sources in the presence and absence of fucoidan. The fucoidan was commercially available fucoidan from *F. vesiculosus* and growth is reported as the maximum optical density (OD_max_) over 48 hours. **(A)** OD_max_ of the four strains (see legend) grown at different concentrations of fucoidan. **(B-C)** OD_max_ of the four strains grown at different concentrations of oligo-alginate (B) or GlcNAc (C) in the presence of 1 mg/mL fucoidan. The shaded area indicates the standard deviation across replicates (*n* = 4). Squares on the right of each panel indicate the average OD_max_ for each strain at the highest concentration of either oligo-alginate (1 mg/mL; B) or GlcNAc (10 mM; C) in the absence of fucoidan (F-). All *p* values are reported in Supplementary Data 1.

Given the strong chemotactic response to fucoidan of the strains tested in spite of their inability to degrade it, we hypothesized that bacteria utilize fucoidan as a signal to navigate to the nutrient-rich surface of macroalgae. To investigate this hypothesis, we tested whether fucoidan inhibits or limits growth in the presence of other carbon sources. We measured the growth of the same four strains at different concentrations of either N-acetylglucosamine (GlcNAc; 0.1–10 mM) or oligo-alginate (0.01–1 mg/mL) in the presence of 1 mg/mL fucoidan (Fig. 4B, C). The two carbon sources are degradation products of chitin and alginate, respectively, two abundant polysaccharides commonly used as nutrient sources by bacteria^65,66^. Although, to our knowledge, the concentrations of GlcNAc and oligo-alginate in the environment have not yet been measured directly, we expect them to reach high values locally within nutrient hotspots, such as macroalgal surfaces and marine particles^67,68^. Three out of four strains exhibited robust growth (OD_max_ = 0.2-0.4) in 1 mg/mL oligo-alginate, without detectable differences between the presence and absence of fucoidan (Fig. 4B; Two-sample t-test, Supplementary Data 1). A similar pattern was observed for GlcNAc: three out of four strains achieved high OD_max_, between 0.25 and 1.5, at elevated substrate concentrations, regardless of the presence of fucoidan (Fig. 4C; Two-sample t-test, Supplementary Data 1). Two strains showed negligible growth with the carbon sources: *Vibrio coralliilyticus* YB1 on 1 mg/mL oligo-alginate with and without fucoidan (OD_max_ = 0.020 and 0.016, respectively; Fig. 4B; two-sample t-test, *p >* 0.05, Supplementary Data 1), and *Pseudocolwellia* sp. G88 on GlcNAc with and without fucoidan (OD_max_ = 0.025 and 0.010, respectively; Fig. 4C; two-sample t-test, *p >* 0.05, Supplementary Data 1). Focusing on *Pseudoalteromonas* sp. ASV16, its growth curves exhibit a similar shape and peak in the presence and absence of fucoidan, for both oligo-alginate and GlcNAc (Fig. S13). Overall, these results indicate that fucoidan did not inhibit bacterial growth on common carbon sources found in fucoidan-rich microenvironments, such as the surface of macroalgae.

## Discussion

We reported the first characterization of the chemotactic response of marine bacteria to the algae-derived polysaccharide fucoidan. As previous studies reported that fucoidan is degraded by very few, specialized strains^23^ and can even have inhibitory effects^33,69,70^, we were surprised to observe strong and widespread chemoattraction toward high fucoidan concentrations (0.1–1 mg/mL), across two types of fucoidans and using two different experimental assays. The magnitude and even the sign of the response were strongly concentration-dependent, prompting us to evaluate whether the tested concentrations are representative of natural conditions. Although such estimates are necessarily approximate and are expected to vary with algal species, environmental conditions, and fucoidan properties, our calculation based on fucoidan release from an incubated *F. vesiculosus* specimen yielded a near-surface concentration on the order of a few mg/mL (estimated at 3.31 mg/mL). This suggests that the concentration range used in our experiments (0.001 to 1 mg/mL) is environmentally plausible for bacteria in the vicinity of algal surfaces. Furthermore, fucoidan is known to exhibit substantial variability in chemical composition and bioactivity across macroalgal species ^28^ and as a function of purification procedures ^71^. To at least partly account for this variability, we tested fucoidan derived from two different macroalgal species, *F. vesiculosus* and *L. hyperborea*, and for different treatments (commercially available, AEX-purified fucoidan and size-fractionated fucoidan): we found that the chemoattraction was largely consistent in all cases, emphasizing the generality of our findings. Several known fucoidan degraders were not tested because they were not motile, consistent with the genomic screen described above (Fig. S1). Since chemotaxis requires motility, these non-motile strains are necessarily unable to exhibit a chemotactic response to fucoidan. One exception is a motile fucoidan degrader/exploiter, *Pseudocolwellia* sp. G88, which was strongly attracted to fucoidan from *L. hyperborea*. Therefore, our findings demonstrate that chemotaxis to fucoidan is a widespread trait among motile marine bacteria, observed across taxonomically diverse lineages spanning multiple families.

Fucoidan is a large molecule (*>*30 kDa) with low diffusivity (*≈* 40 *µ*m^2^ s*^−^*^1^) and is therefore expected to generate steep chemical gradients^72^. Such gradients are likely to persist longer than those of most low–molecular-weight metabolites because fucoidan undergoes limited microbial degradation and is continuously released from macroalgae. This persistence suggests that fucoidan may act as a complementary cue relative to small, rapidly diffusing metabolites, potentially providing a more temporally stable signal for locating algal surfaces over longer timescales. Chemotaxis research to date has overwhelmingly focused on small molecules^38^ and only recently have there been efforts to quantify bacterial responses to polysaccharides^39,40^. As a consequence, the mechanisms by which chemotactic bacteria detect and respond to gradients of macromolecules remain unknown. Here we showed that steep fucoidan gradients elicit strong bacterial chemotactic behavior. Using two different experimental platforms (the ISCA and the gradient generator), we experimentally and numerically confirmed that fucoidan gradients remained steep for relatively long times, spanning the duration of our experiments (60 min for the ISCA; 10 min for the gradient generator). In the gradient generator, where the temporal evolution of the gradient could be precisely resolved by imaging fluorescent fucoidan, bacterial taxis occurred predominantly during periods when the fucoidan gradients were steepest, before falling below a threshold (0.005 mg*/*mL *µ*m*^−^*^1^). In the environment fucoidan is continuously exuded and accumulates at high concentrations within the diffusive boundary layer of brown macroalgae^9^, which will result in steep gradients persisting over prolonged times. Under these conditions, bacteria with chemotactic abilities similar to those measured here will be able to exquisitely sense the fucoidan gradient and rapidly navigate toward algal surfaces. We thus propose that the chemotactic behaviors we observed will also occur in natural marine environments, primarily in the vicinity of the surface of macroalgae. This close-range mode of action contrasts with that of smaller, more rapidly diffusing chemoattractants such as amino acids (Fig. 3D), which are expected to form shallower gradients extending further from the algal surface^73^. We therefore posit that bacterial navigation toward macroalgae may involve cues acting over different spatial scales: longer-range attraction to small metabolites, followed by fucoidan acting as a closer-range cue that reinforces and potentially maintains bacterial accumulation once bacteria are already near the algal surface. Distinguishing the relative contributions of these short- and long-range cues to bacterial navigation in situ is an important direction for future work. Finally, we note that, under strong flow, such as that driven by waves in shallow waters, we expect the extent of chemotaxis to decrease, as both the extent of the diffusive gradient and the ability of bacteria to navigate it via motility will deteriorate.

The strong chemotaxis of marine bacteria to fucoidan suggests that large polysaccharides are detected by a currently unknown mechanism, different from the well-described signal cascade induced by smaller molecules. The most common chemotactic signal transduction mechanism is through transmembrane chemoreceptors, either by binding directly to ligands or through binding proteins^52^. Although other polysaccharides can be sensed through periplasmic ligand-binding proteins, the large size of fucoidan (*>*30 kDa) makes periplasmic transport difficult, implying low sensing capability^74^. Furthermore, we consider it highly unlikely that all tested strains have evolved a specific ligand-binding protein for fucoidan, especially since only a subset of the strains was isolated from fucoidan-rich environments. Nutrient influx detection, an alternative chemotaxis mechanism without transmembrane receptors^52,75^, is equally unlikely, as it would require the breakdown of this recalcitrant molecule. We suggest that the chemotaxis response to fucoidan may be driven by alternative forms of signal transduction that do not involve ligand-receptor binding and constitute a physiological response of the cell, such as those reported for pH, oxygen, and osmolarity^52^. Our observations thus call for further work to understand this sensing mechanism.

Interestingly, a subset of the strains tested exhibited chemorepulsion at low fucoidan concentrations (0.01 – 0.001 mg/mL). Some of these strains (*Pseudoalteromonas* sp. ASV16, *Pseudoalteromonas* sp. AS80, *Pseudoalteromonas* sp. AS81 and *Pseudocolwellia* sp. G88) thus exhibited a biphasic response to fucoidan from *F. vesiculosus*, combining attraction at high concentration with repulsion at low concentration. Similar biphasic chemotactic responses have been reported in *E. coli* ^58,76^, where they have been interpreted as adaptive strategies allowing bacteria to navigate efficiently in heterogeneous chemical environments. The repulsion we observed was generally weaker and less prevalent than the attraction. No repulsion toward fucoidan from *L. hyperborea* was detected. Repulsion toward fucoidan from *F. vesiculosus* occurred in only three of the 14 strains tested and resulted in *I_C_* values as low as 0.3, corresponding to a 70% reduction in cell accumulation relative to the control. In contrast, attraction occurred in all 14 strains and reached *I_C_* values as high as 800, corresponding to an 800-fold increase in cell accumulation.

After AEX purification of *F. vesiculosus* fucoidan, the repulsive response was no longer generally observed, and only one strain showed significant repulsion, indicating that compounds responsible for repulsion were removed or substantially reduced in concentration by the purification process. Additionally, because fucoidan’s low diffusivity generates a concentration gradient that decays outward from the algal surface, the low fucoidan concentrations associated with repulsion may only occur over a narrow spatial region at a distance from the algal surface^72^. These differences suggest that chemorepulsion is unlikely to dominate bacterial responses to fucoidan in natural environments. Nevertheless, the ecological significance of weak repulsion at low concentrations combined with strong attraction at high concentrations remains unclear and will require further investigation to assess its prevalence, robustness under natural conditions, and ecological role. Understanding the underpinning of the chemorepulsion response we detected at low fucoidan concentrations in some strains from fucoidan-rich environments warrants future work. Testing additional specialist degrader and non-degrader motile isolates would help resolve whether the underlying catabolic capabilities of the strains align with chemorepulsion at low fucoidan concentrations.

Fucoidan is a recalcitrant molecule due to its complex chemical structure^24^, with degradation restricted to a limited number of specialist bacteria that possess dedicated enzymatic repertoires^77,78^. In line with this, none of the strains we tested grew on fucoidan as the sole carbon source. The strong anionic properties of fucoidan have also been associated with antibacterial activity in various non-marine bacteria^32,79,80^. In our experiments with marine strains, we observed neither bactericidal nor bacteriostatic activity of fucoidan in the presence of the carbon sources tested. Recent work has suggested that brown macroalgae use the biochemical composition of their surface to exert a strong selective pressure on their microbiome^13,81^. Our results indicate a strong attraction of marine bacteria to an important metabolite of macroalgae across many different bacterial species, regardless of whether they were isolated from fucoidan-rich environments or not. Thus, it seems unlikely that fucoidan alone is responsible for this strong selective pressure on the macroalgae microbiome. Instead, it seems more plausible that such selective pressure results from the combined action of fucoidan with other macroalgal metabolites, many of which have antimicrobial properties (e.g., phlorotannins, sterols, halogenated terpenes) ^82,83,84^. Future work should aim to disentangle the relative contributions of fucoidan and other macroalgal compounds in shaping the microbial communities surrounding macroalgae.

Our findings reveal widespread chemotaxis toward a large macromolecule that most marine bacteria cannot directly use as a growth substrate. Several small metabolites, such as amino acids and dimethylsulfoniopropionate (DMSP), have been identified as infochemicals that are not necessarily degraded, but instead mediate interactions between environmental bacteria and hosts, including plants, phytoplankton, and corals^85,86^. These results support the view that chemotaxis is not restricted to small nutrient substrates^39^, but instead also serves as a behavioral strategy to locate favorable environments^41^. Indeed, a recent analysis indicates that more than one quarter of 926 experimentally identified compounds do not support growth, but function solely as chemotactic signals^38^. Our findings provide the first demonstration that a large molecule such as fucoidan can also serve as an infochemical (i.e., a chemical cue that is not a growth substrate) for bacteria in the ocean. Under this framework, the physiological benefit of chemotaxis to fucoidan does not derive from its consumption: rather than requiring a direct growth payoff, chemotaxis to fucoidan can be favored if it reliably guides bacteria toward macroalgal surfaces and similar microenvironments enriched in other, readily usable carbon sources, as we discuss below.

Based on our results, we propose a new ecological model in which chemotaxis toward fucoidan promotes interactions between marine bacteria and macroalgae. Fucoidan is released at high concentrations from the surfaces of macroalgae (in the order of 1 mg/mL), where its low diffusivity and limited biodegradability generate steep and persistent chemical gradients. Therefore, within this close-range zone, these gradients constitute a highly reliable chemical signal that bacteria can utilize to navigate to macroalgal surfaces. Many motile bacteria are capable of sensing these gradients and thus can chemotax toward and accumulate on the surfaces of macroalgae, where they gain access to environments rich in labile nutrients^87^. Hence, chemotaxis to fucoidan can facilitate microbial recruitment to macroalgal surfaces. Our strain panel, however, was dominated by non-specialist taxa, several of which belong to genera that can have algicidal effects, rather than bacterial groups that are commonly recognised as beneficial macroalgal symbioses (e.g., *Sulfitobacter*, *Hyphomonas*, *Rhizobium* ^88,89^). Nevertheless, our genomic screen (Fig. S1) shows that several potential symbiotic taxa (e.g., *Sulfitobacter*, *Hoeflea*, *Roseibium*) possess complete flagellar biosynthesis and chemotaxis gene complements, representing promising candidates for future testing. Elucidating the extent and specificity of these reciprocal interactions will be essential for a comprehensive understanding of the role of fucoidan in shaping macroalgal holobionts.

Beyond macroalgal surfaces, fucoidan released into the water column may also accumulate within organic matter particles^20^, where polysaccharides produced by both macro and microalgae are known to concentrate during particle formation and aging^90^. Although fucoidan concentrations on marine particles remain unknown, steep fucoidan gradients associated with organic-rich marine particles could similarly act as chemotactic cues, potentially contributing to their colonization. While fucoidan itself is largely recalcitrant and unlikely to represent a major direct carbon source for the bacteria studied here, chemotaxis toward it is still relevant to carbon cycling, by concentrating motile bacteria at sites of organic matter turnover where other, more labile compounds are locally available for consumption and remineralization. In summary, this work identifies fucoidan as a potent and widespread chemotactic signal in the marine environment. We thus propose that chemotaxis toward fucoidan is an important mechanism structuring bacterial interactions with macroalgae. Ultimately, our findings highlight how macromolecular chemical landscapes can shape microbial behavior, with potential implications for carbon cycling in the ocean.

## Methods

### Bacterial cell culture

The bacterial strains used in the experiments were selected from previously characterized chemotactic isolates ^39,23,91,92,48^. Individual strains were cultured in Marine Broth 2216 (BD Difco), either diluted 1:10 or used at full strength in artificial seawater (Instant Ocean, Spectrum Brands), according to strain-specific requirements (Supplementary Table 1). Cultures were incubated overnight at 27 *^◦^*C with shaking at 180 rpm for 16 to 24 h, according to strain-specific requirements (Supplementary Table 1).

### ISCA experiments with fucoidan

ISCA chips were manufactured following established protocols ^43,44^. Each ISCA consisted of an array of 20 wells (5 × 4), each of a volume of 110 *µ*L and featuring an 800 *µ*m-diameter inlet port that connected the well to the external environment. This configuration enabled the diffusion of the solutes into the surrounding medium, forming microscale gradients representative of those occurring near the algal surface. Before each experiment, chemoattractant solutions were prepared by dissolving the appropriate amount of fucoidan powder from *F. vesiculosus* (Sigma-Aldrich) or *L. hyperborea* in sterile, 0.2 *µ*m-filtered and autoclaved artificial seawater (ASW; Instant Ocean, Spectrum Brands) to achieve the desired final concentrations. ASW alone served as a negative control. All solutions were then filtered through 0.2 *µ*m syringe filters (Millipore) to remove particles and potential contaminants. The chemoattractant or control solutions were loaded into individual wells using 1 mL sterile syringes (Codan) fitted with 27G needles (Henke Sass Wolf). Prior to each ISCA experiment, the motility of the overnight bacterial culture was confirmed by phase contrast microscopy. Cell concentrations were then determined by flow cytometry (CytoFLEX, Beckman Coulter) and diluted to 10^6^ cells/mL in ASW. ISCAs were incubated for 1 h in 50 mL of the bacterial solution. Following incubation, the contents of each well were individually retrieved using sterile syringes and transferred to 1.5 mL Eppendorf tubes for downstream analysis. Samples were stained with SYBR Green I (1:10,000 final dilution; Thermo Fisher Scientific), incubated in the dark for 15 min, and analyzed using a CytoFLEX S flow cytometer (Beckman Coulter). Forward scatter (FSC), side scatter (SSC), and green fluorescence (FITC channel) were recorded. Bacterial population was identified based on SSC and SYBR Green fluorescence (Fig. S14), and absolute cell abundances were calculated from a standardized acquisition volume of 30 *µ*L per sample. The chemotactic index (*I_C_*) was calculated as the ratio of the number of cells detected in the chemoattractant-containing wells to those of the control wells containing only ASW. Statistical analysis was performed using ANOVA and Dunnett’s post-hoc test on log-transformed cell counts. To account for multiple testing across strains within each heatmap, Dunnett-adjusted p-values were further adjusted across strains separately for each concentration using the Benjamini–Hochberg procedure. Raw flow cytometry data were exported as CSV files, and heatmaps and plots were generated using Python v3.9 and the Seaborn visualization package.

### Chemotaxis assay with the microfluidic gradient generator

The gradient generator microfluidic devices were manufactured using standard single-layer soft lithography as previously described ^62,60^. Polydimethylsiloxane (PDMS; Sylgard 184, Dow Corning) was cast onto SU-8 molds, cured, and bonded to glass microscope slides (76 mm *×* 26 mm) via oxygen plasma treatment. Before each experiment, the devices were pre-conditioned by flushing with 0.5% (w/v) bovine serum albumin (BSA; Sigma-Aldrich) in artificial seawater (ASW; Instant Ocean, Spectrum Brands) to minimize bacterial adhesion to channel surfaces. Fucoidan solutions were prepared by dissolving the appropriate amount of fucoidan powder from *F. vesiculosus* or *L. hyperborea* in ASW. Bacterial suspensions were prepared by washing overnight cultures three times with ASW and resuspending the cells to a final concentration of 10^8^ cells/mL. The three inlets were loaded with (i) fucoidan in ASW (top inlet), (ii) bacterial suspension in ASW (center inlet), and (iii) ASW only (bottom inlet). Solutions were introduced using a syringe pump (Harvard Apparatus), with syringe sizes adjusted to maintain a 4:1:4 flow rate ratio across the three streams. Flow was maintained for at least 2 minutes to establish laminar stratification, after which it was stopped to allow the chemoattractant gradient to develop via diffusion. The chemotactic response of the bacterial population was then recorded. Imaging was performed using phase-contrast microscopy (10×, 0.13 NA objective; Nikon Ti-E) at 1 frame per second for a total duration of 10 min, using a sCMOS camera (Zyla, Andor Technology). Technical replicates were obtained by reinitiating the flow for 2 min to reset initial conditions, followed by cessation of flow and renewed image acquisition. A fresh microfluidic device was used for each of the three biological replicates. Cell positions were extracted from the recordings using previously developed analytical scripts^45^ implemented in MATLAB (MathWorks). The observed chemotactic response was quantified using the accumulation index *β* ^93,94,62^, defined as

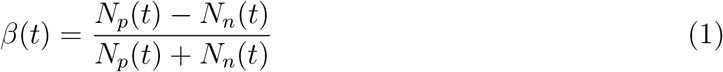

where *N_p_*(*t*) and *N_n_*(*t*) represent the instantaneous number of cells at time *t* within predefined positive and negative accumulation regions ^62^, respectively, with the difference normalized by the total number of cells in these two regions.

### Growth assays with fucoidan and carbon sources

The bacterial strains were precultured in Marine Broth 2216 (BD Difco), undiluted (100%) or diluted 1:10 in artificial seawater depending on their optimal growth conditions, and incubated at 27 *^◦^*C for 18 h until reaching the stationary phase. Growth assays were performed in complete F/2 medium (Bigelow Laboratory), supplemented with 1 mg/mL of commercially available fucoidan from *F. vesiculosus* (Sigma-Aldrich) and varying concentrations of oligo-alginate or N-acetylglucosamine (GlcNAc; Sigma-Aldrich) as additional carbon sources. Oligo-alginate was generated by enzymatic depolymerization of 2% (w/v) sodium alginate using 1 U/mL alginate lyase (Sigma-Aldrich) at 37 *^◦^*C for 24 h, following established protocols^91^. All media and substrate solutions were sterile-filtered through 0.2 *µ*m syringe filters (Millipore) before use. Overnight cultures were diluted into the assay medium to an initial optical density at 600 nm (OD_600_) of 0.001. Growth experiments were carried out in 96-well microplates with 200 *µ*L medium per well, using four technical replicates per condition. Plates were incubated at 27 *^◦^*C, and growth dynamics were monitored by measuring OD_600_ every 10 min over 48 h using a microplate reader (BioTek Instruments) controlled by Gen5 software. Growth curves were processed and analyzed using Python v3.9, and data visualization and handling were performed with the Pandas library.

### AEX purification and desalting of fucoidan

Crude fucoidan from *F. vesiculosus* (85% purity) (Marinova) and crude *L. hyperborea* fucoidan extracted from the environment were further purified using a precipitation sequence with CaCl2, followed by anion exchange chromatography^95^, and a precipitation and desalting method using ethanol^96^. In brief, a 5 g/L fucoidan solution was treated with 2 M CaCl2 in an alkaline environment created with NaOH. The solution was centrifuged to separate precipitated impurities from dissolved fucoidan in the supernatant. For subsequent anion exchange chromatography, the fucoidan solution was neutralized and diluted in running buffer A (0.5 M NaCl, 20 mM Tris-HCl) and loaded onto an ANX-HiTrap column. After washing with buffer A, the purified anionic polysaccharide was eluted with running buffer B (4 M NaCl, 20 mM Tris-HCl) in 12 mL per starting 1 mg crude fucoidan. To desalt the samples, the extracted fucoidan in the eluents was precipitated using 70-80% EtOH, filtered over a combusted GF/F filter, and washed with 70-80% EtOH. The white solid was reconstituted from the filter in ultra-pure water, then freeze-dried for long-term preservation.

### Size-fractionation of fucoidan

Fractionation of AEX-purified and commercially available (Sigma) fucoidan from *Fucus vesiculosus* was performed using Amicon Ultra centrifugal filter units (Sigma) with a molecular-weight cutoff of 5 kDa. Filter membranes were first equilibrated by adding 10 mL of artificial seawater (ASW) and centrifuging for 10 min at 4000 rpm; the flow-through was discarded. The equilibrated membrane was then loaded with 10 mL of the fucoidan solution and centrifuged for 20 min at 4000 rpm; The flow-through, corresponding to the fraction *<*5 kDa (LMW), was collected and stored at *−*20 *^◦^*C until use. The retained material (HMW; *>*5 kDa) was washed twice by adding 10 mL of ASW to the filter unit and centrifuging for 20 min at 4000 rpm, collecting the flow-through after each wash. The washed retentate was then resuspended in 10 mL of ASW by gentle pipetting directly onto the filter membrane, ensuring complete recovery of the high-molecular-weight fraction, and stored at *−*20 *^◦^*C until use.

### Live-dead assay with propidium iodide in microfluidics growth chambers

Microfluidic devices for single-cell viability assays were fabricated and operated as in previous works ^97^. Briefly, the device consisted of an array of 100 square microchambers (50 µm × 50 µm × 1 µm) located between two parallel main channels (100 µm width, 20 µm height). The microchambers allowed for spatial confinement of individual bacterial cells while maintaining perfusion via the main channels. *Pseudoalteromonas* sp. ASV16 cultures were introduced into the device by injecting 0.1 mL of cell suspension with 10^8^ cells/mL into one of the main channels with a 200 *µ*L pipette. Cells entered the microchambers via diffusion and passive trapping. After loading, a constant flow of artificial seawater (ASW) supplemented with 1 mg/mL AEX-purified fucoidan from *F. vesiculosus* was applied within the main channels at a rate of 5 mL/hour to maintain environmental stability and exposure to nutrient over the duration of the test. At the end of the incubation period, propidium iodide solution (PI; 10 *µ*g mL*^−^*^1^) was introduced into the device and allowed to incubate for 15 min to label membrane-compromised cells (non-viable). Live and dead cells were imaged directly within the microfluidic chip using a Nikon Ti inverted fluorescence microscope equipped with a 40x objective and mCherry filter. The images were then analyzed to quantify cells using Fiji/ImageJ and plots were produced using Python v3.9.

### Live-dead assay with propidium iodide in bulk

The viability of *Pseudoalteromonas* sp. ASV16 was assessed using propidium iodide (PI) staining following exposure to different concentrations of AEX-purified fucoidan from *F. vesiculosus*. Overnight bacterial cultures in 10% 2216 Marine Broth were diluted 1:1000 in artificial seawater (ASW) and exposed to fucoidan at final concentrations of 1 mg/mL or 0.001 mg/mL. Control conditions included a heat-killed positive control and two negative controls. For the positive control, bacterial suspensions were incubated at 80 *^◦^*C for 15 min, then cooled on ice for 5 min. Negative control 1 consisted of cells stained with SYBR Green I only (no PI), while negative control 2 consisted of cells stained with both dyes in ASW without fucoidan. All conditions were prepared in triplicate and the samples were collected and analyzed 1, 2, 4, 6, and 24 hours after treatment. Staining was performed on a total volume of 500 *µ*L. PI (stock concentration 1 mg/mL) was added at a 1:1000 dilution, and SYBR Green I (1:100 working solution) was added to achieve a final 1:10,000 dilution. The samples were incubated for 15 min at 4 *^◦^*C in the dark. Flow cytometry analysis was conducted on a CytoFLEX S (Beckman Coulter). The samples were run at a flow rate of 30 *µ*L min*^−^*^1^ for 60 s per sample. Fluorescence data were acquired using CytExpert software, and gating was applied to distinguish SYBR Green^+^ (total bacteria) and PI^+^ (membrane-compromised) populations. The fraction of dead cells was quantified using the following equation

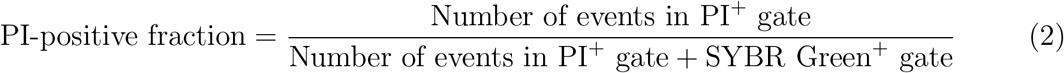

Data were analyzed and plotted using the Seaborn library in Python v3.9.

### Preparation of fluorescently labelled fucoidan

A starting 5 mg/mL of AEX-purified fucoidan solution from *F. vesiculosus* was cooled in an ice bath and adjusted to pH 8. A diethylammonium phosphate (DAMP, 2.5 M) solution was added to buffer the system during the reaction. With the addition of a cyanodiazolyl phosphate (CDAP) solution (100 mg/mL in Acetonitrile), the reaction was started and ran for 15 min. The reaction mixture was then directly loaded on a self-packed SEPHADEX G-50 column with a borate buffer (0.2 M) as mobile phase. For the column chromatography, an AKTA Start system was used with published methods^98^. The eluted activated sugar was collected and 4 mg fluorescein amine added. The solution was allowed to react overnight and subsequently purified with another column chromatography using a SEPHADEX G-50 based column and a phosphate buffer (50 mM, with 100 mM NaCl) as mobile phase. The solvent of the eluted fluorescently labelled fucoidan was exchanged for ultra-pure water using ultrafiltration tubes (molecular-weight cut off at 5 kDa). The fluorescently labelled fucoidan was concentrated in that step and stored at room temperature.

### Simulations of fucoidan and amino acid diffusion

Solute diffusion was simulated numerically in COMSOL Multiphysics assuming purely diffusive transport according to

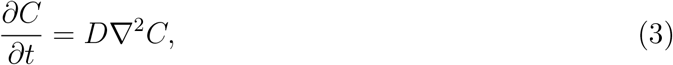

To simulate diffusion in the 3-inlet chip, we considered a 1 mm^2^ square channel with periodic boundary conditions along the horizontal (*x*) direction and no-flux boundary conditions along the vertical (*y*) direction, representing a section of the main channel. Solute was initially confined to the upper 400 *µ*m of the channel at a normalized concentration *C*_0_ = 1. Simulations were run for 10 min, and the concentration profile along the *y* axis was extracted at the midpoint of the channel (*x* = 500 *µ*m). For the ISCA, we modeled the concentration field along the axis extending from the bottom of the well to the external medium through the well outlet. The ISCA well was initially filled with solute at concentration *C*_0_ = 1, while the surrounding medium was solute-free. Concentration profiles were expressed as *C/C*_0_ and plotted as a function of distance from the well outlet, where 0 mm denotes the outer border of the outlet and *−*3 mm the bottom of the well. Profiles were evaluated from 0 to 7200 s to capture gradient evolution during 2 h incubation. For fucoidan and amino acids, we used an approximated diffusion coefficient of 40 *µ*m^2^ s*^−^*^1^ and 400 *µ*m^2^ s*^−^*^1^, respectively. Concentration values are reported normalized by the initial concentration, *C*_0_.

### Genetic engineering of *Vibrio coralliilyticus*

A chemotaxis-deficient mutant of *Vibrio coralliilyticus* YB1 was created by deleting the *cheA1* gene. The deletion of the *cheA1* gene encoding protein (EEX33918.1) was performed using homologous recombination using a mutagenesis plasmid. To create the plasmid, the primers B450-cheA-up-EcoRI-F and B50-cheA1-up-OEX-R was used to amplify approximately 800 bp of DNA upstream of *cheA1* via PCR, similarly the primers B450-cheA1-down-OEX-F and B450-cheA1-down-XbaI-R were used to amplify approximately 800 bp downstream of *cheA1*. The two products were purified and used as products for overlap extension PCR^99^ together with primers B450-cheA1-up-EcoRI-F and B450-cheA1-down-XbaI-R. The approximately 16 kb product was purified and then digested with the restriction enzymes EcoRI and XbaI. The product was then cloned into the suicide plasmid pSW4426T^100^ digested with the same restriction enzymes to create the plasmid pBU260. To create the knockout deletion, a previously described protocol using homologous recombination was used^101^. Briefly, the mutagenesis plasmid pBU260 in *E. coli beta* 3914^100^ was introduced into *V. coralliilyticus* YB1 via conjugation. After two successful rounds of recombination, potential mutants were initially screened using soft-agar chemotaxis assays^101^ followed by PCR using the primers B450-cheA-up-EcoRI-F and B450-cheA1-down-XbaI-R. All confirmed *cheA1* mutants were screened for any growth defects by comparing growth over time against the parental strain (YB1).

### Dynamic light scattering measurements

To estimate the hydrodynamic radius (Rh) and calculate the diffusion coefficient (D) of commercially available fucoidan from *F. vesiculosus*, dynamic light scattering (DLS) measurements were performed using a 3D-DLS instrument in cross-correlation geometry (LS Instruments). Scattering data were collected at a backscattering angle of 140°. Each measurement consisted of at least six consecutive runs of 1 min. The intensity time-correlation function was evaluated using cumulant analysis in the instrument software. Fucoidan concentrations of 2.5, 5 and 10 mg/mL were analyzed. All measurements were performed at 23°C using artificial seawater as the buffer.

### Rheometer measurements of fucoidan viscosity

Viscosity measurements were performed to assess whether fucoidan solutions for commercially available fucoidan from *F. vesiculosus* were different from control seawater, using a stress-controlled rheometer (Kinexus Lab+, Netzsch) equipped with a cone-plate geometry (40 mm diameter, 1° angle). Samples were equilibrated at 25 *^◦^*C until stable prior to measurement. A solvent trap was used to minimize desiccation during the measurement procedure. Flow curves were obtained by applying a shear rate sweep from 0.001 s*^−^*^1^ to 100 s*^−^*^1^ in logarithmic steps. Apparent viscosity was calculated as the ratio of shear stress to shear rate. All measurements were performed in triplicate, and results were averaged. Data analysis and visualization were conducted using the Seaborn library (Python v3.13).

### Tracking of bacterial motile behavior in the presence of fucoidan

To characterize the effect of different concentrations of commercially available fucoidan from *F. vesiculosus* on bacterial motile behavior, overnight cultures of *Pseudoalteromonas* sp. ASV16 were grown in 10% Marine Broth 2216 (BD Difco), diluted 1:500 in artificial seawater (Instant Ocean, Spectrum Brands), and exposed to six treatment conditions: (i) 10 mg/mL fucoidan, (ii) 1 mg/mL fucoidan, (iii) 0.1 mg/mL fucoidan, (iv) 0.01 mg/mL fucoidan, (v) 0.001 mg/mL fucoidan, and (vi) artificial seawater only (negative control). After 15 min incubation at room temperature, 30 *µ*L of each cell suspension was placed at the center of a custom imaging chamber composed of a standard microscope slide and coverslip, separated by approximately 1 mm of silicone rubber. Motility was recorded using phase contrast microscopy (Nikon Ti-E) with a 20×, 0.45 NA objective. Videos were acquired at 30 frames per second for 30 s using a Hamamatsu CMOS camera, with an image resolution of 2044 *×* 2048 pixels (0.326 *µ*m per pixel). Cell tracking was performed as previously reported^39^. Background noise was removed by subtracting the median projection of each image stack. Cells were tracked using TrackPy v0.6.4^102^. Trajectories were analyzed in Python v3.9, and data visualization was performed using the Pandas and Seaborn libraries

### Genomic screening for motility and chemotaxis machinery

To evaluate the genetic capacity for motility and chemotaxis across members of a fucoidan-associated bacterial community, we screened 29 published genome assemblies from this community^48^ against 152 unique motility-associated KEGG Orthology (KO) identifiers (Kanehisa & Goto, KEGG database). These KOs spanned structural (flagellar biosynthesis: *fli*, *flg*, *mot*), regulatory (e.g. *flhA/B*, *flhDC*, *fleQ/fleS/fleR*), and chemosensory (*che* genes, methyl-accepting chemotaxis proteins, type IV pilus/twitching motility genes) categories, enabling comparative profiling of behavioral and locomotive traits across community members. KO hits per genome were tabulated and visualized as a clustered heatmap.

### NMR measurements of different fucoidan preparations

Freeze dried samples were dissolved in 0.5 mL D2O. The 1H-NMR-Spectra were measured with an NMR-Spectrometer Bruker Avance NEO 600 at 600.50 MHz. Measurements were performed at 25 °C and with 64 scans. The chemical shifts (*δ*) are reported in parts per million (ppm) and deuterated water was used as reference (4.79 ppm). Spectra were compared and annotated according to^95^ and a measured 1H-NMR of L-fucose.

## Data availability

All data supporting the findings of this study are available within the Article and its Supplementary Information.

## Supporting information

Supplementary Data for the main manuscript

Supplementary Data for the supplementary information

## Acknowledgments

We thank members of the Stocker laboratory and PriME community for helpful discussions and feedback on the manuscript. We thank Russell Naisbit for scientific editing. We thank Estelle Clerc for providing bacterial strains used in this study. This work was supported by the Swiss National Science Foundation (grant number 207488) and by the Simons Foundation (PriME grant number 542395FY22).

## Contributions

V.A., J.B.R., J.K., J.H.H. and R.S. conceived the work and designed the experiments. V.A., R.J.H., J.K., S.C., M.R., and L.B. performed experiments, with help from L.B., B.U., M.B., A.S., and G.S.U; V.A., J.K., R.F., R.J.H., M.R., and A.S. analysed experimental data. R.F. performed simulations. All authors discussed and interpreted the results. V.A., J.B.R., J.K., R.J.H., and R.S. wrote the manuscript with input from all authors.

