## Supplementary Information for "Widespread attraction of marine bacteria to the macroalgae-derived polysaccharide fucoidan"

### Supplementary Information for
### Widespread attraction of marine bacteria to the macroalgae-derived polysaccharide fucoidan

Valerio Anelli[1], Jean-Baptiste Raina[2], Riccardo Foffi[1], Linda Biehler[3], Richard J. Henshaw[1], Blake Ushijima[4], Margot Bligh[3], Giovanni Stefano Ugolini[1], Sam Charlton[1], Milad Radiom[5], Natasha Blitvic[6], Raffaele Mezzenga[5], Andreas Sichert[7], Jan-Hendrik Hehemann[2,3], Johannes Keegstra*[1,8], Roman Stocker*[1]

[1]Institute for Environmental Engineering, ETH Zurich, Switzerland
[2]Max Planck Institute for Marine Microbiology, Bremen, Germany
[3]Center for Marine Environmental Sciences, University of Bremen, Germany
[4]Department of Marine Biology, UNC Wilmington, USA
[5]Department of Health Sciences and Technology, ETH Zurich, Switzerland
[6]School of Mathematical Sciences, Queen Mary University of London, UK
[7]Institute of Molecular Systems Biology, ETH Zurich, Switzerland
[8]School of Biological and Behavioral Sciences, Queen Mary University of London, UK

This PDF includes:
Supplementary Notes 1 and 2
Supplementary Figures S1–S14
Supplementary Table 1

#### Supplementary Notes

##### Supplementary Note 1

To determine an ecologically relevant concentration of fucoidan for the experiments, we estimated the concentration expected at the algal surface ($C_s$) using Fick's law of diffusion for a spherical geometry[1]:

$$Q = 4\pi D(C_s - C_0)\, k. \tag{1}$$

The parameters were calculated as follows:

- The leakage rate ($Q$) was estimated by incubating a thallus of *F. vesiculosus* in 1 L of Milli-Q water for 1 h. After incubation, the released mucilage was collected both from the surrounding water and by rinsing the algal surface to recover slime remaining attached to the thallus. Fucoidan was then purified from the collected liquid by liquid chromatography, yielding approximately 6 mg of fucoidan after 1 h, corresponding to an estimated leakage rate of $\sim 6\ \mathrm{mg\,h^{-1}}$. Because the amount recovered varied by species, season, and collection site, this value should be considered an order-of-magnitude estimate.

- The diffusivity coefficient ($D$) for fucoidan was estimated using the Stokes-Einstein equation, with the hydrodynamic radius ($R_h$) of *F. vesiculosus* determined by 3D dynamic light scattering (3D-DLS) (Fig. S8) and averaged across three different concentrations (2.5, 5, and 10 mg/mL) as $R_h = 5.5$ nm. This yielded a value of $D = 36.4\ \mu\mathrm{m}^2\,\mathrm{s}^{-1}$.

- The fucoidan concentration in bulk water ($C_0$) was measured as 0.006 $\mathrm{g\,m^{-3}}$[2]. Since this is much smaller than the concentration close to the algae, this was approximated as zero for simplicity.
- The capacitance ($k$) is a geometric factor that describes diffusive transport from an object into the surrounding fluid. In the case of a sphere, the diffusive capacitance is equal to its radius ($k = R$). To obtain an order-of-magnitude estimate for the macroalga, we approximated it as an equivalent sphere with the same total surface area. We measured a dry biomass of approximately 15 g and, assuming a water content of 80%[3], estimated a wet biomass of approximately 75 g. Using a specific surface area of 0.2 $\mathrm{m^2\,g^{-1}}$, this corresponds to a total algal surface area of $A \approx 15$ $\mathrm{m^2}$. Using the relation $A = 4\pi R^2$, this surface area corresponds to an equivalent spherical radius of $R = \sqrt{A/(4\pi)} \approx 1$ m. We therefore used $k \approx 1$ m. Because real macroalgae have highly irregular morphologies rather than spherical shapes, this approximation should be interpreted as a simplified geometric estimate.

Rearranging Eq. 1, we obtain:

$$C_s = \frac{Q}{4\pi D R} \approx 3.31\ \mathrm{mg\,mL^{-1}}$$

This estimate suggests that fucoidan concentration experienced by bacteria near the algal surface is on the order of 1 mg/mL, once the uncertainty associated with the $k = R$ geometric approximation is taken into account. We therefore used 1 mg/mL as the highest concentration in the experiments.

#### Supplementary Note 2

To confirm that the observed gradient characteristics were not specific to the microfluidic setup, we simulated fucoidan diffusion in the ISCA using COMSOL (Fig. S11), where gradients are steady rather than pulsed and persist for the 1-hour incubation period. Consistent with the microfluidic measurements and COMSOL simulations shown in Fig. 3D, simulations of the ISCA geometry confirmed the formation of a steep chemical gradient emanating from the well (Fig. S11). Fucoidan diffusion remained confined to the immediate vicinity of the well throughout the 1-hour incubation period (0-0.5 mm), with negligible fucoidan concentrations detected at distances of 1 mm from the well outlet (Fig. S11), indicating that both experimental platforms generate similarly steep fucoidan gradients. Together, these findings confirm that the bacterial chemotactic response is specifically elicited by steep fucoidan gradients.

### Supplementary Figures

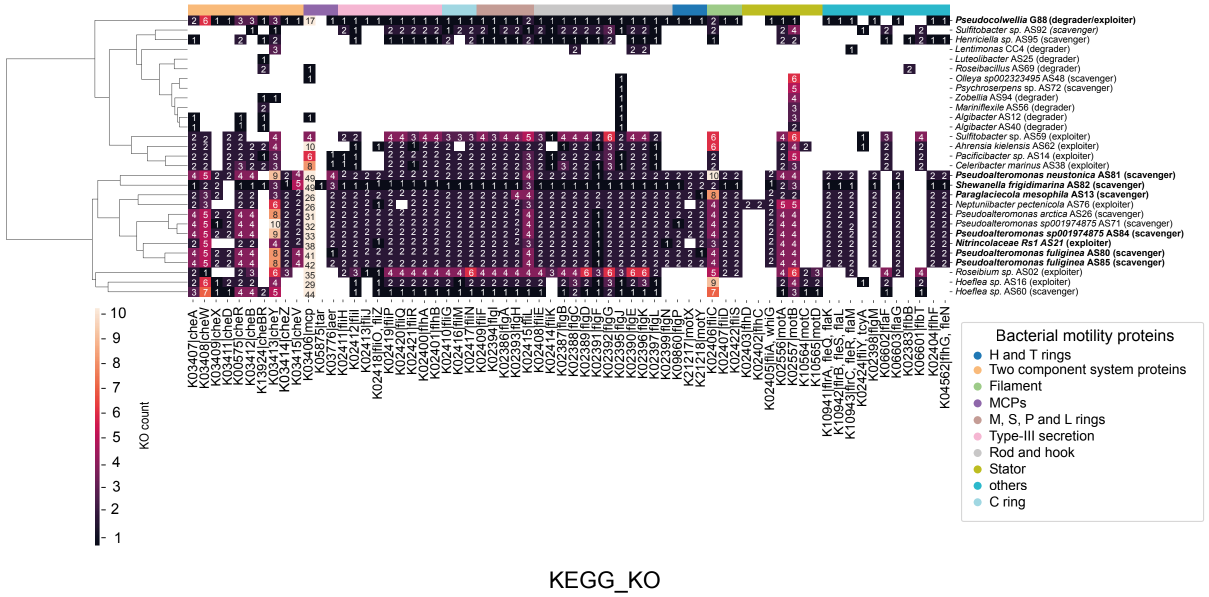


Figure S1: **Genomic screen for motility and chemotaxis machinery across a fucoidan-associated bacterial community**. Heatmap showing counts of KEGG Orthology (KO) hits for 152 motility- and chemotaxis-associated gene families across 29 bacterial genomes from a fucoidan-degrading community[4]. Genomes are hierarchically clustered based on KO profile similarity (dendrogram, left). Each strain is annotated with its proposed ecological role within the fucoidan degradation pathway — degrader (initial breakdown of intact fucoidan), scavenger (uptake of degradation products released into the environment), or exploiter (opportunistic use of breakdown products without contributing to degradation) — following the division-of-labor framework of Sichert et al. Columns are grouped and color-coded by functional gene category (top color bar; legend, right): M/S/P/L-ring, MCP, type-III secretion, two-component system, rod and hook, H/T-ring, stator, filament, C-ring, and others. Cell color and overlaid values indicate KO count per gene per genome (scale, bottom left); blank cells indicate no hit. Strains with a near-complete complement of flagellar biosynthesis and chemotaxis genes (lower clade) correspond to the taxa used in chemotaxis assays in this study and are highlighted in bold.

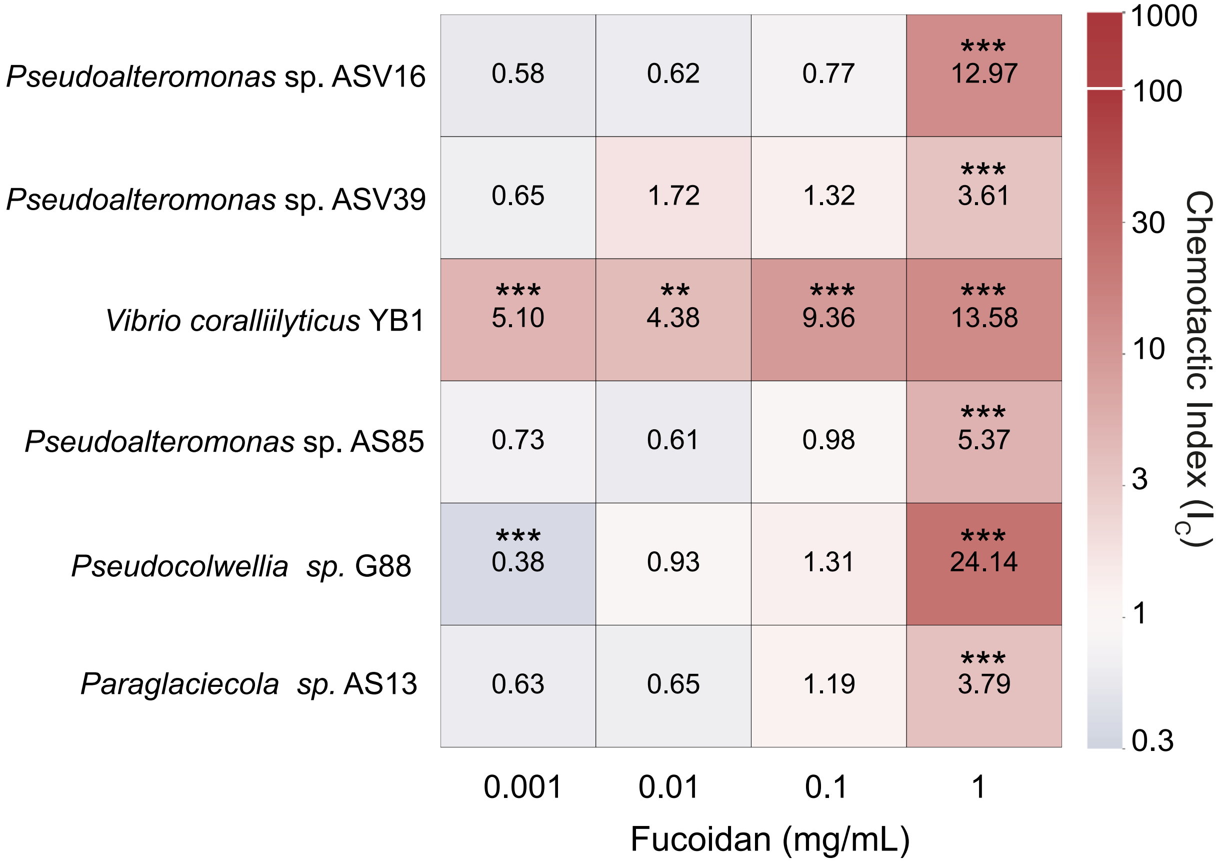


Figure S2: **ISCA screening of bacterial chemotactic response towards AEX-purified fucoidan from *F. vesiculosus*.** Heatmap summarizing the chemotactic responses of 6 bacterial strains towards 4 different ecologically-relevant concentrations of AEX-purified fucoidan from *F. vesiculosus*. Each row represents the average chemotactic index $I_C$ for a different bacterial strain ($n = 3$), and each column represents a different fucoidan concentration. Stars indicate chemotactic responses significantly higher (attraction) or lower (repulsion) than the ASW control (Dunnett's test, with Benjamini–Hochberg $p$-values adjustment for testing across multiple strains $p < 0.05$, $**p < 0.01$, $***p < 0.001$). All $p$-values are reported in Supplementary Data 2.

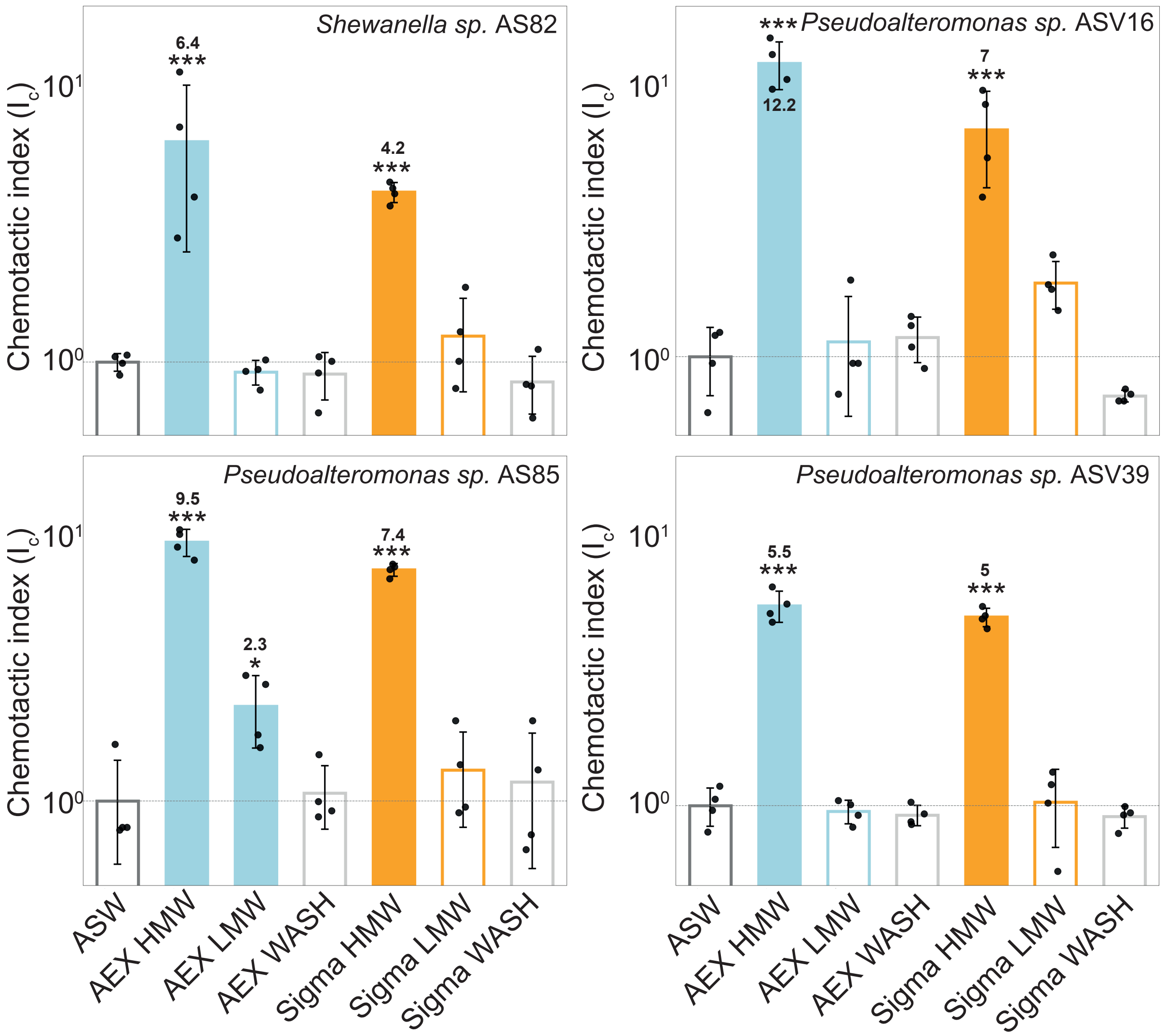


Figure S3: **ISCA screening of bacterial chemotactic response towards size fractionated AEX-purified and commercially available fucoidan from *F. vesiculosus*.** Chemotactic response of *Pseudoalteromonas* sp. ASV16, *Pseudoalteromonas* sp. ASV39, *Pseudoalteromonas* sp. AS85 and *Shewanella* sp. AS82 to 1 mg/mL of the high-molecular-weight fraction (HMW; > 5 kDa) and low-molecular-weight fraction (LMW; < 5 kDa), respectively, following size-fractionation of AEX-purified (AEX) and commercially available (SIGMA) fucoidan, including the washing step from the fractionation process as a control. The horizontal dashed gray line indicates the control $I_C = 1$ value of Artificial Seawater (ASW). Stars and filled barplots indicate chemotactic responses significantly higher (attraction) than to the control performed with ASW (* = $p < 0.05$; ** = $p < 0.01$; *** = $p < 0.001$; Dunnett's test, with Benjamini–Hochberg p-value adjustment for testing across multiple strains) and corresponding average $I_C$ values reported immediately above the bar. All $p$ values are reported in Supplementary Data 2.

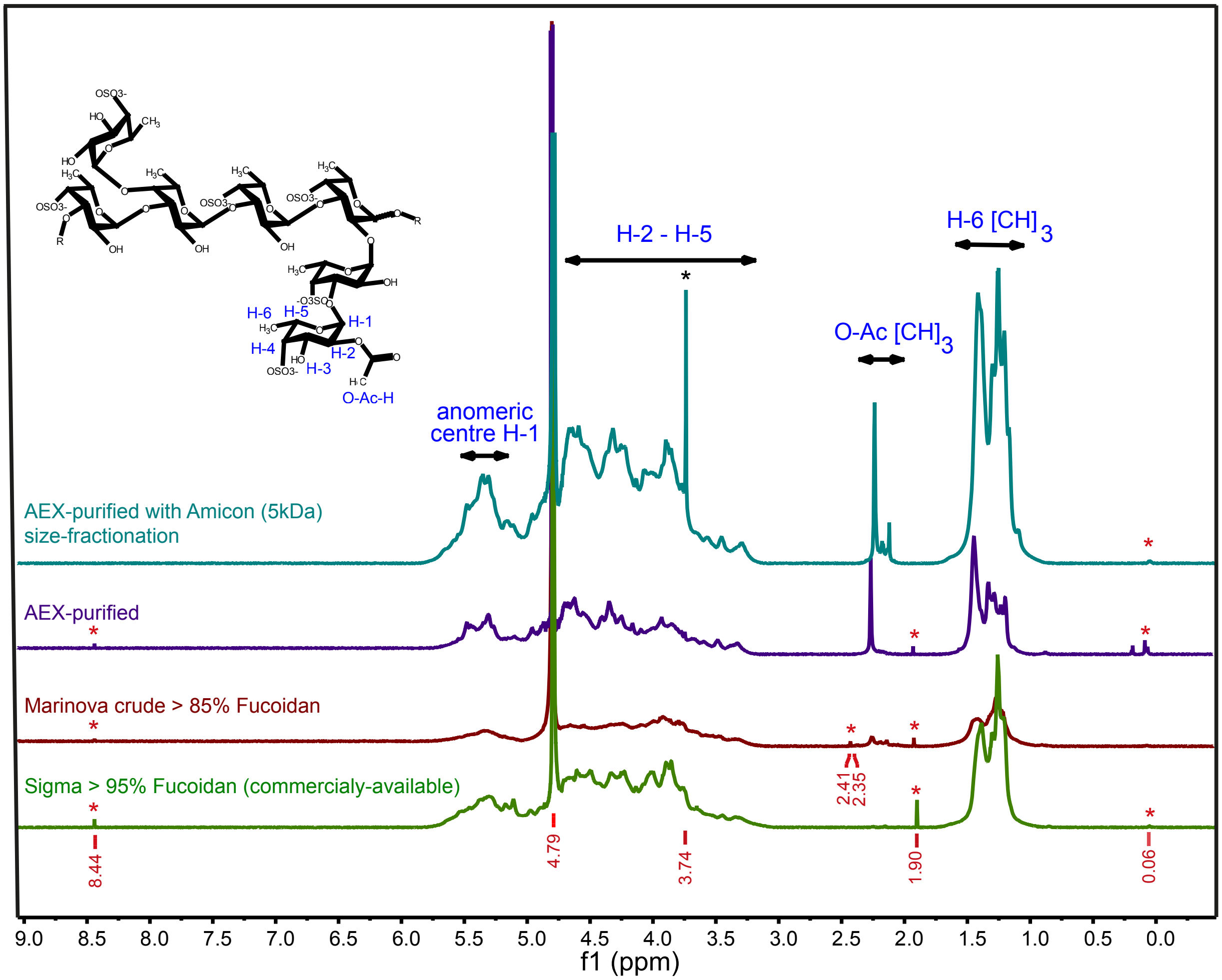


Figure S4: **$^1$H NMR spectra of fucoidan from *Fucus vesiculosus* at different stages of purification** $^1$H NMR spectra (DO, 600.15 MHz, 64 scans) of fucoidan preparations at four stages of purification, from top to bottom: AEX-purified fucoidan following Amicon (5 kDa) size-fractionation; AEX-purified fucoidan before size-fractionation; crude fucoidan (Marinova, >85% fucoidan); and commercially available fucoidan (Sigma-Aldrich, >95% fucoidan). A proposed chemical structure of fucoidan is shown in the top-left inset, with proton assignments corresponding to the labeled spectral regions (blue) indicated directly on the structure: the anomeric proton (H-1), the ring protons (H-2–H-5), the O-acetyl proton (O-Ac-H), and the methyl protons (H-6, [CH]). Asterisks (*) denote external single peaks not belonging to the fucoidan structure; corresponding chemical shifts are annotated below the spectra. The peak at 4.79 ppm is annotated as $D_2O$, the solvent used to resuspend the different fucoidan preparations, while the peak at 3.74 ppm indicates Tris from the elution buffer.

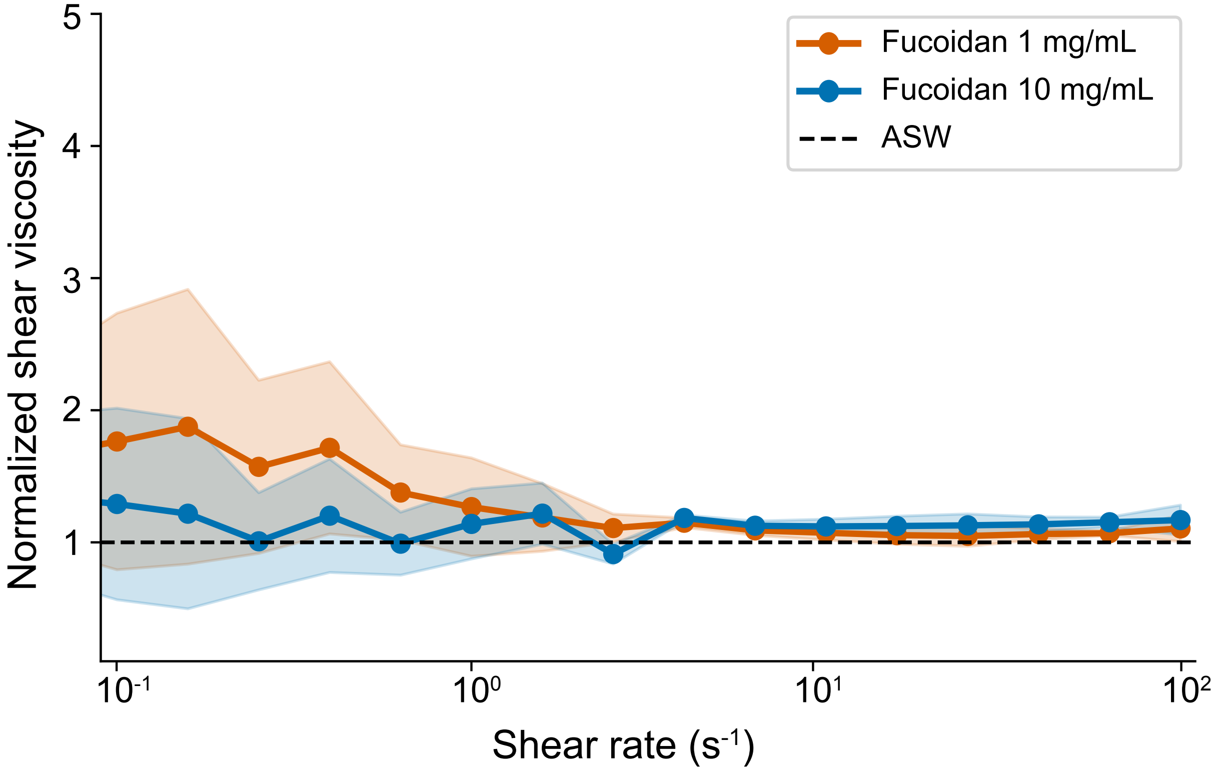


Figure S5: **Rheometer measurements of fucoidan viscosity**. Shear viscosity of fucoidan solutions (1 and 10 mg/mL) measured across a range of shear rates and normalized to the viscosity of artificial seawater (ASW). Shaded regions indicate propagated standard deviations from replicate measurements of fucoidan solutions and ASW ($n = 3$). The dashed horizontal line marks the ASW reference (normalized viscosity $= 1$). Across the tested shear rates, fucoidan solutions exhibited viscosities comparable to ASW. The average shear viscosity was, respectively, 0.0065 $\mathrm{Pa \cdot s}$ for 1 mg/mL fucoidan, 0.0088 $\mathrm{Pa \cdot s}$ for 10 mg/mL fucoidan, and 0.0055 $\mathrm{Pa \cdot s}$ for ASW. Statistical analysis was performed using a two-sample $t$-test. All $p$-values are reported in Supplementary Data 2.

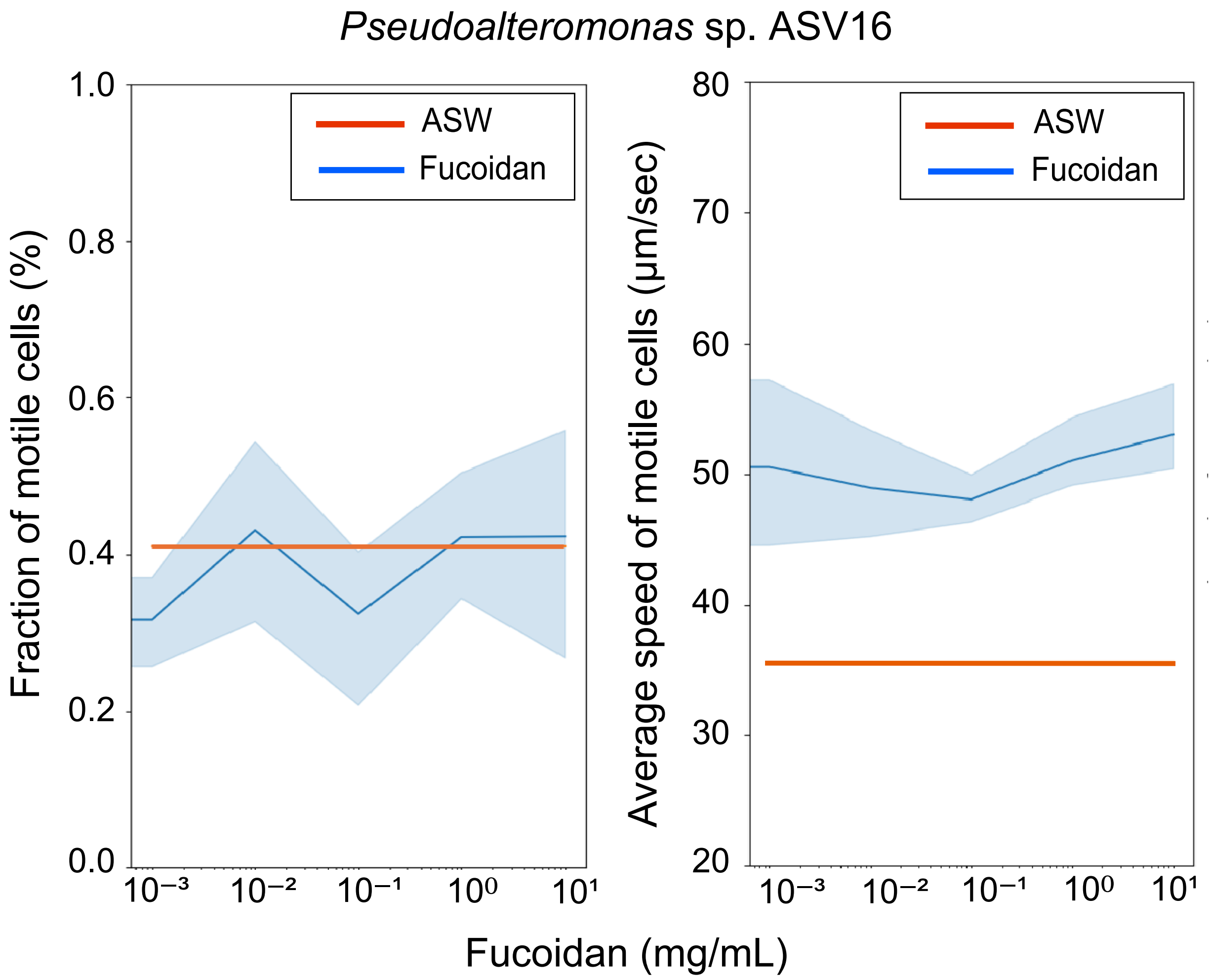


Figure S6: **Motile behaviour of *Pseudoalteromonas* sp. ASV16 across a range of fucoidan concentrations**. Fraction of motile cells (left) and average swimming speed (right) of *Pseudoalteromonas* sp. ASV16 measured in uniform concentrations of fucoidan from *Fucus vesiculosus* across a range of concentrations (0.001 to 10 mg/mL). Blue lines indicate measurements in the presence of fucoidan, with shaded regions representing standard deviations across replicates ($n = 3$). Orange lines indicate the control condition in artificial seawater (ASW).

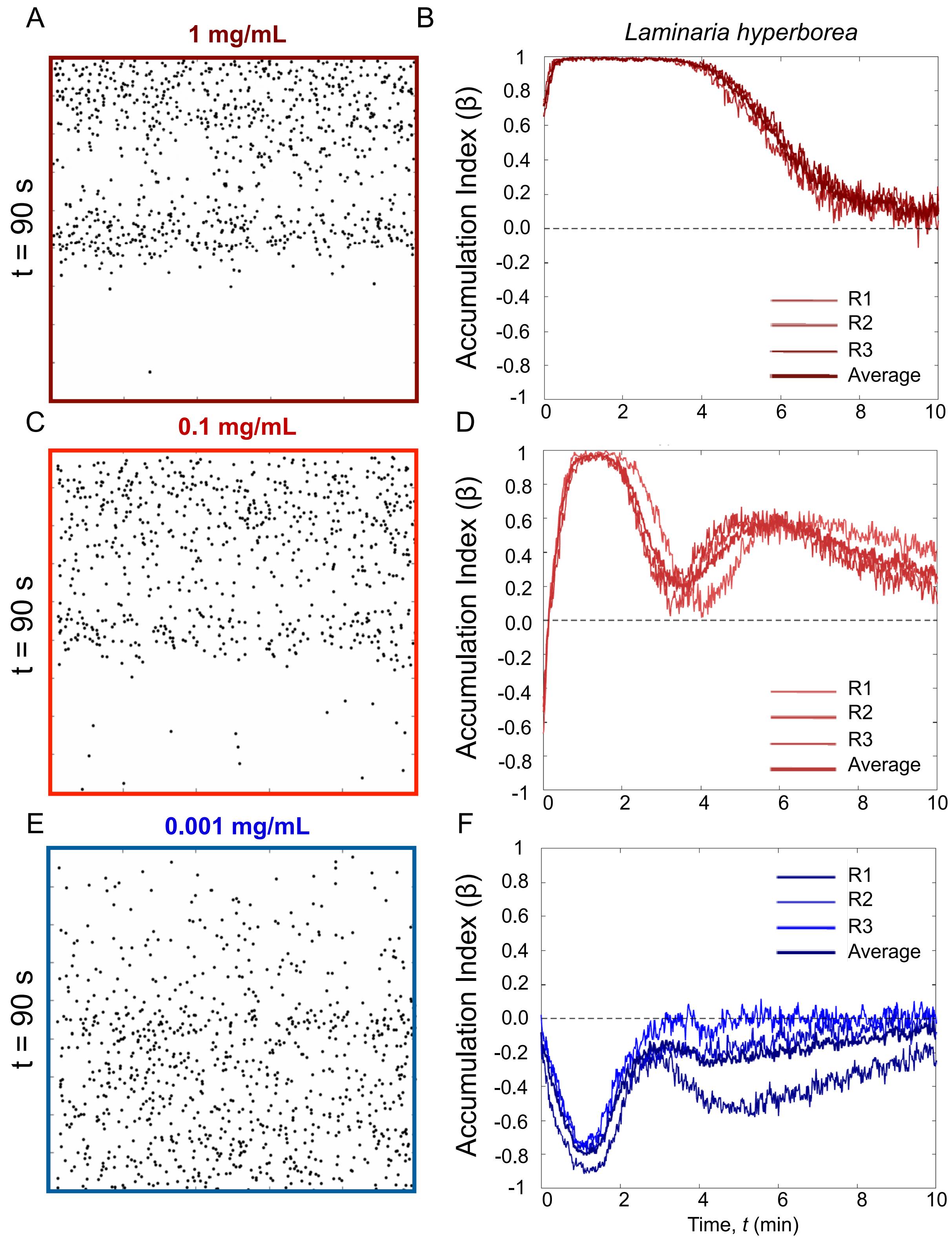


Figure S7: (Caption on next page)

Figure S7: **Chemotactic response of *Pseudoalteromonas* sp. ASV16 to fucoidan from *L. hyperborea* in the gradient generator**. **A, C, E** Snapshots of bacterial cell positions in the microfluidic channel 90 s after stopping the flow for high (1 mg/mL), intermediate (0.1 mg/mL), and low (0.001 mg/mL) fucoidan concentrations, respectively. Each black dot represents a single bacterial cell. **B, D, F** Time series of the bacterial accumulation index $\beta(t)$ for the corresponding fucoidan concentrations. Lines show three independent technical replicates (R1–R3), with the thicker line indicating the average response.

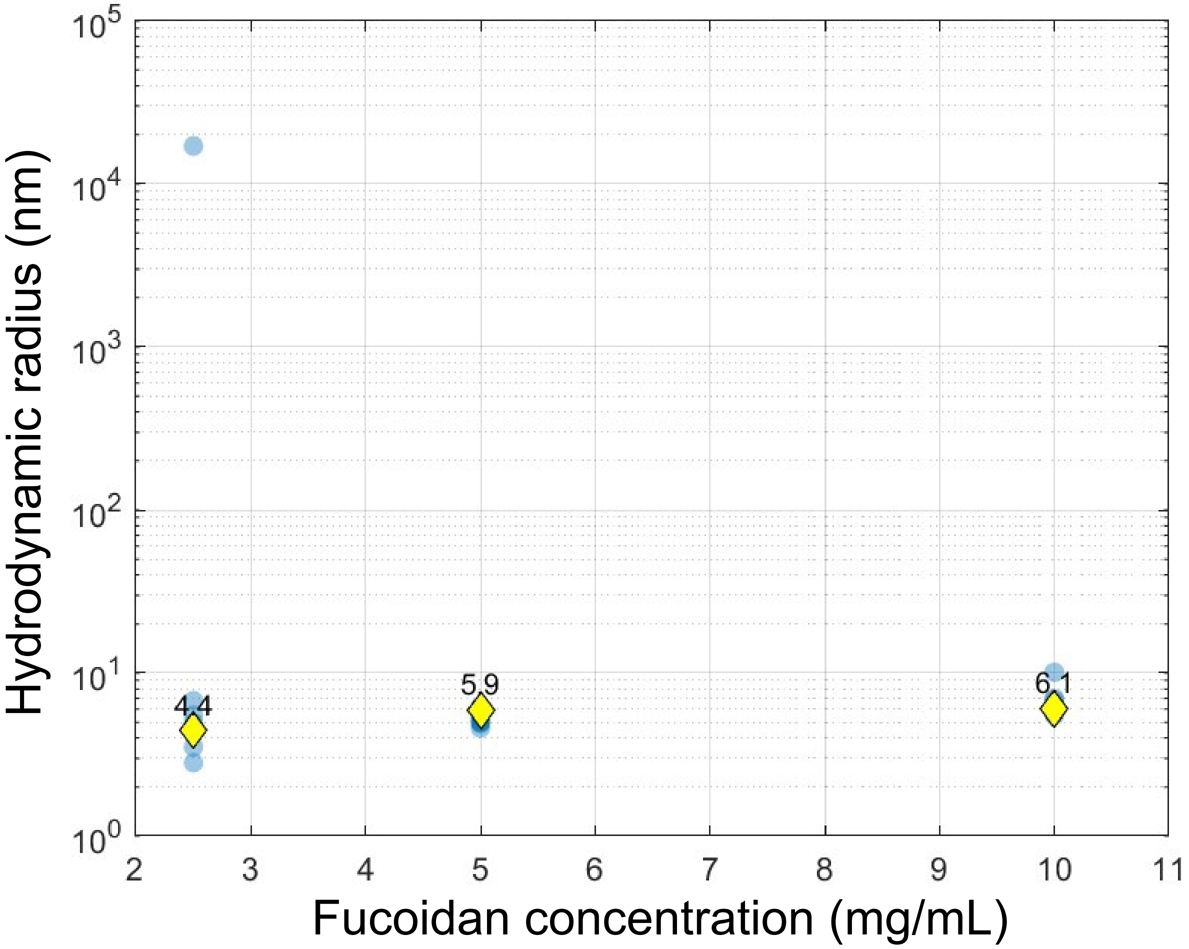


Figure S8: **Dynamic light scattering (DLS) characterization of commercially available fucoidan from *F. vesiculosus***. Hydrodynamic radius of fucoidan measured across three concentrations: 2.5, 5, and 10 mg/mL. The yellow diamond indicates the peak (modal) value for each concentration. Blue circles represent individual replicates ($n = 12$).

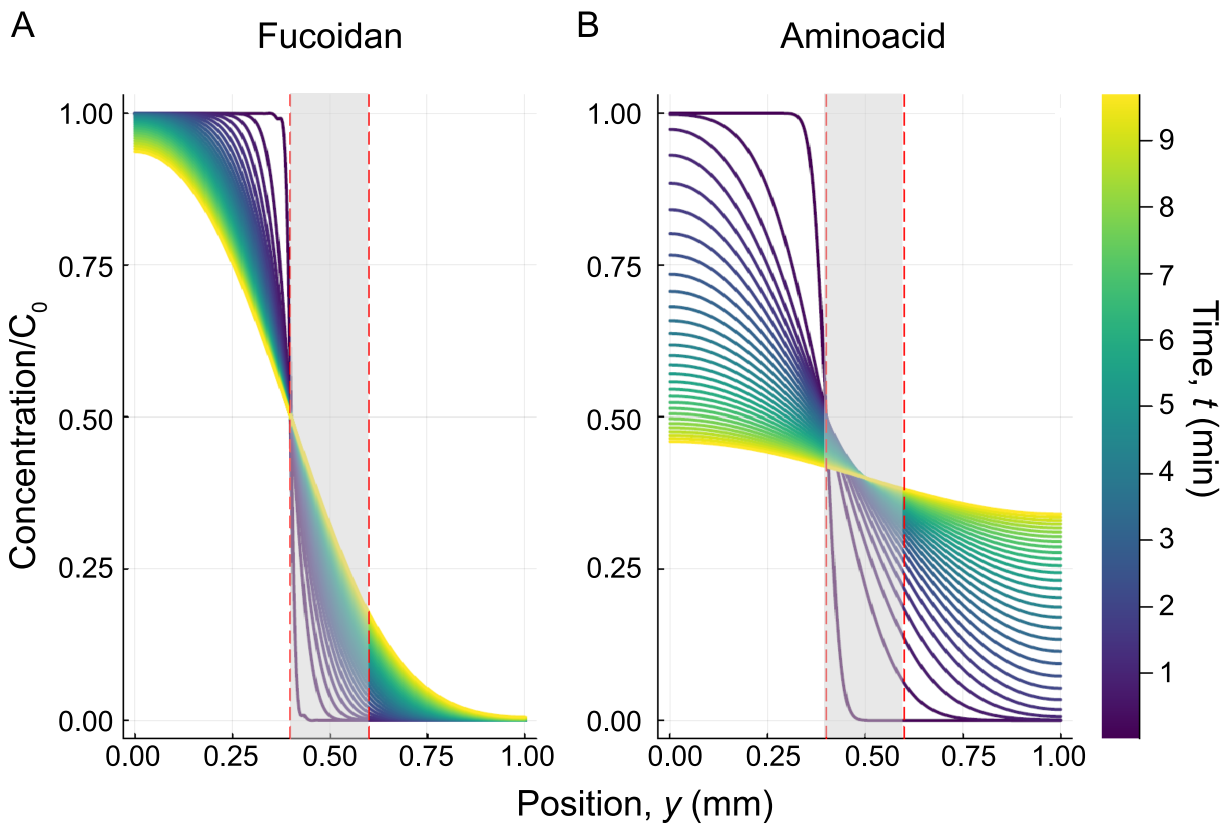


Figure S9: **Simulation of the evolution of concentration profiles for fucoidan and amino acids in the 3-inlet gradient generator**. **(A)** Fucoidan. **(B)** Amino acid. The vertical axis shows concentration normalized by the initial concentration value ($C_0$). The horizontal axis shows the 1 mm width of the chip from top to bottom, with the chemical source located at the top. Dashed red lines and the gray area indicate the band where bacteria were localized at time 0. Each curve represents a different time point at 20-s intervals. For fucoidan and amino acids, we used an approximated diffusion coefficient of $40\,\mu\text{m}^2\,\text{s}^{-1}$ and $400\,\mu\text{m}^2\,\text{s}^{-1}$.

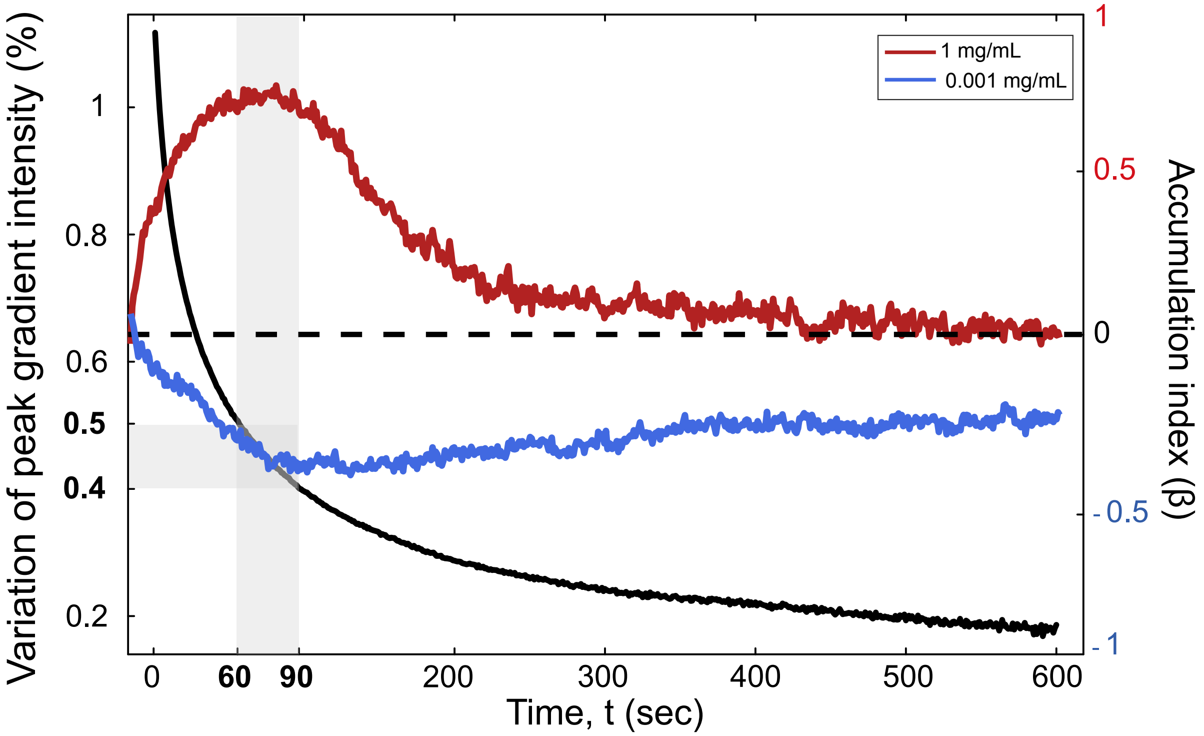


Figure S10: **Variation of fucoidan peak concentration gradient in the microfluidic chip compared to bacterial response**. The black curve shows the peak concentration gradient in the 3-inlet gradient generator as a function of time, estimated by fluorescence intensity. Units are defined such that 1 a.u. (1% or 0.01 $\mathrm{mg\,mL^{-1}\,\mu m^{-1}}$) corresponds to the fluorescence level associated with the input fucoidan concentration. Red and blue curves indicate the $\beta(t)$ index for the bacterial response to high (1 mg/mL) and low (0.001 mg/mL) concentrations, respectively, of AEX-purified fucoidan from *F. vesiculosus*. Gray areas indicate the time window and peak-gradient intensity threshold required for bacteria to respond to the gradient.

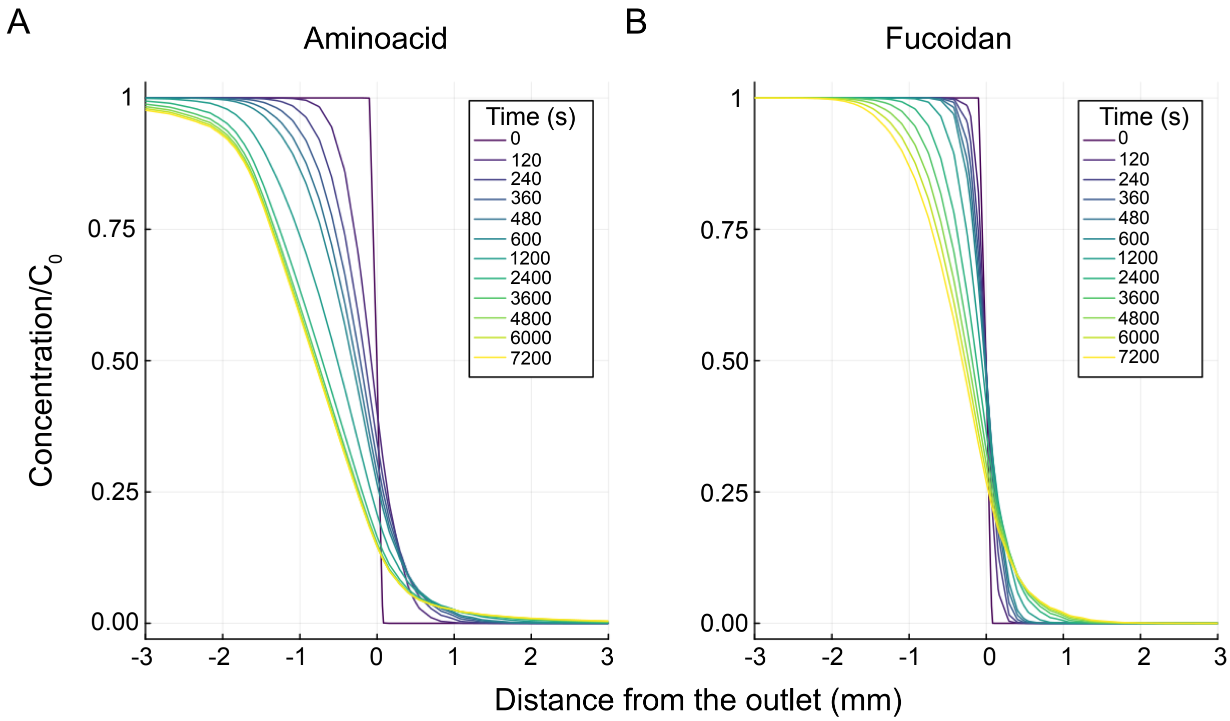


Figure S11: **Simulation of the evolution of concentration profiles for fucoidan and amino acids in the ISCA**. Curves show the concentration of amino acid **(A)** and fucoidan **(B)** normalized by the initial concentration ($C_0$) as a function of distance from the ISCA well outlet, where 0 mm represents the outer border of the outlet and $-3$ mm the bottom of the well, where the gradient originates. Each curve represents a different time point over 0-2 h of incubation (color scale). For fucoidan and amino acids, we used an approximated diffusion coefficient of respectively $40\,\mu\mathrm{m}^2\,\mathrm{s}^{-1}$ and $400\,\mu\mathrm{m}^2\,\mathrm{s}^{-1}$.

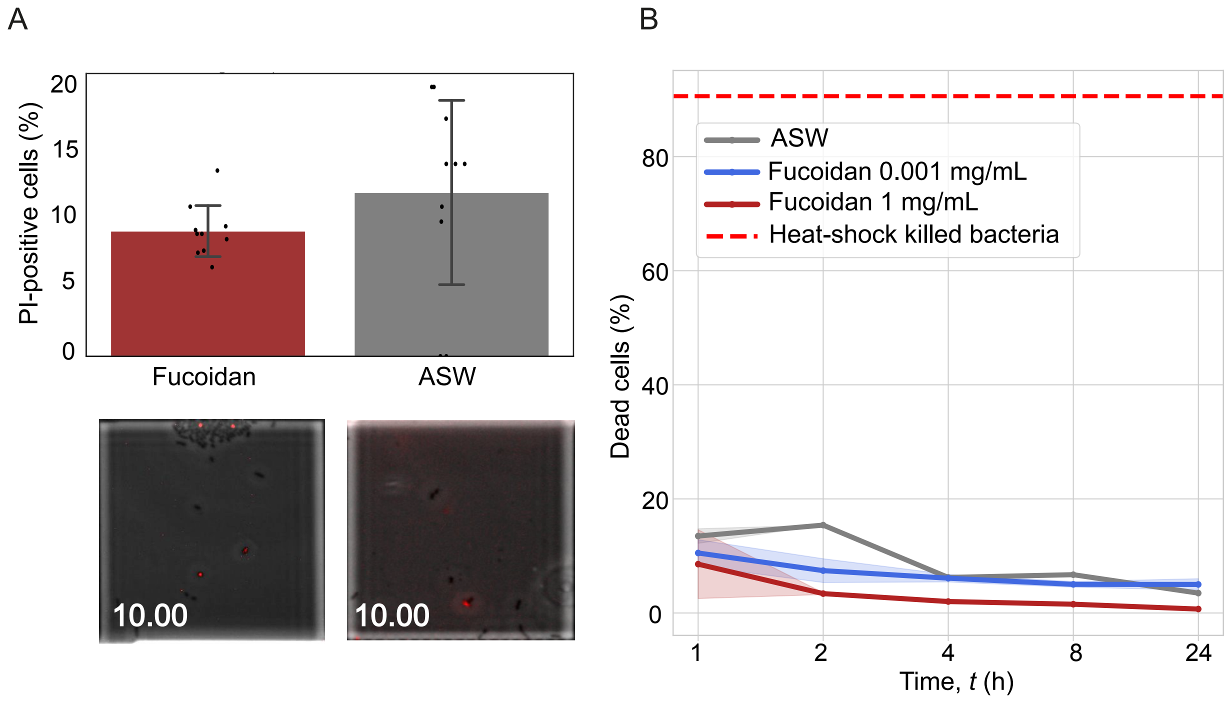


Figure S12: **Live/dead assays with *Pseudoalteromonas* sp. ASV16 and commercially available fucoidan from *F. vesiculosus*.** **(A)** The top plot shows the percentage of dead cells in the microfluidic growth chamber for bacteria exposed to a high concentration of fucoidan (1 mg/mL; red) and the ASW control (gray). Dead bacteria were identified by propidium iodide (PI) staining. Quantification was performed by counting PI-positive cells in different chambers at the 10 h time point, and error bars indicate the standard deviation ($n = 10$). Bottom images show representative snapshots of the growth chambers under the two conditions at the time point used for quantification ($t = 10$ h). **(B)** Flow-cytometry quantification of the percentage of dead cells in bulk experiments for high (1 mg/mL) and low (0.001 mg/mL) fucoidan concentrations, compared to the negative ASW control (gray) and a positive heat-killed bacterial control (dashed red line). The shaded region indicates the standard deviation across replicates ($n = 3$). Statistical analysis was performed using a two-sample $t$-test. All $p$-values are reported in Supplementary Data 2.

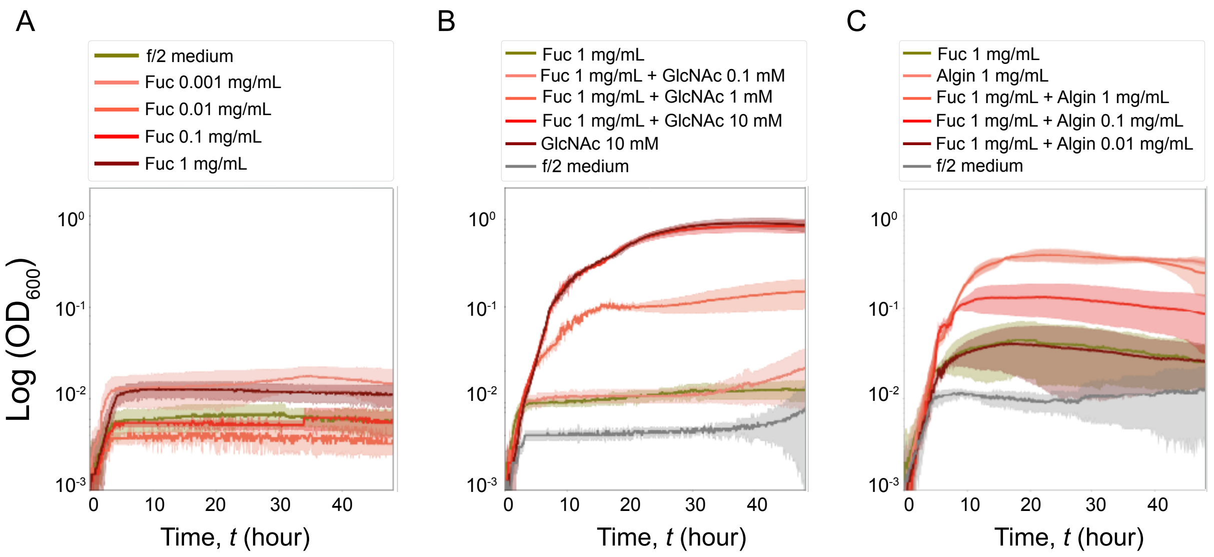


Figure S13: **Growth of *Pseudoalteromonas* sp. ASV16 in the presence of commercially available fucoidan from *F. vesiculosus* and additional carbon sources**. **(A)** Growth curves of *Pseudoalteromonas* sp. ASV16 showing log(OD) over the 48 h experiment in response to different concentrations of fucoidan (Fuc), compared with the f/2 medium control. **(B)** Growth curves showing log(OD) over 48 h in response to different concentrations of N-acetylglucosamine (GlcNAc) in the presence of a high background concentration of fucoidan (1 mg/mL), compared with the f/2 medium control (gray). **(C)** Growth curves showing log(OD) over 48 h in response to different concentrations of oligo-alginate (Algin) in the presence of a high background concentration of fucoidan (1 mg/mL), compared with the f/2 medium control (gray). Shaded areas indicate the standard deviation across replicates ($n = 4$).

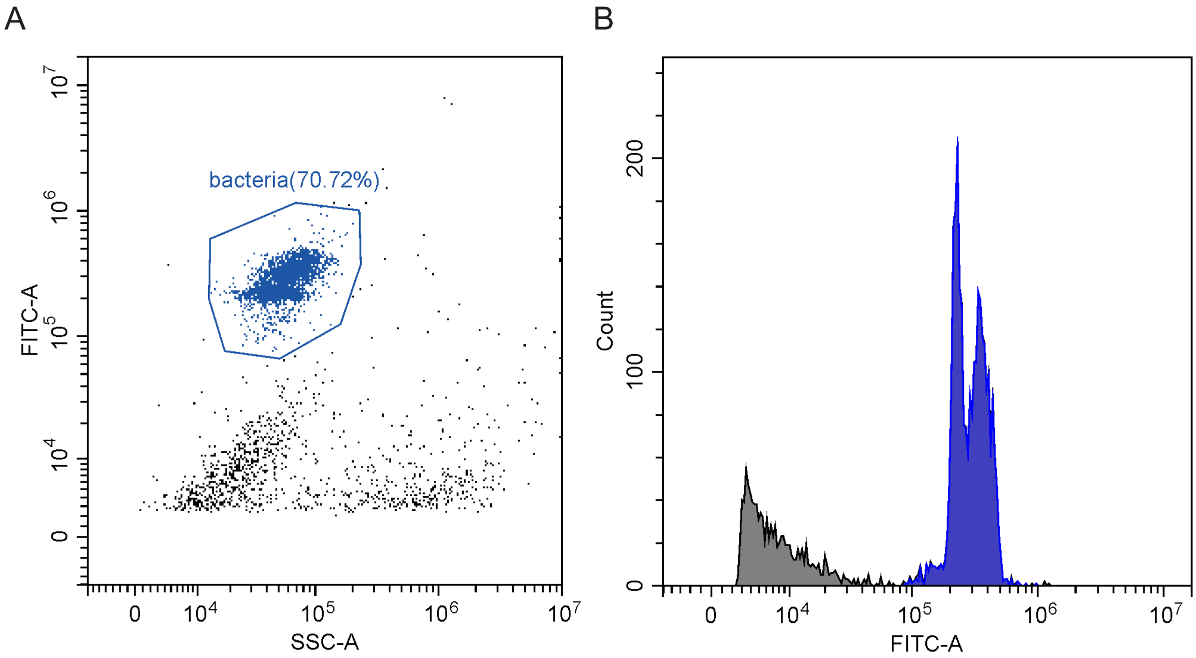


Figure S14: **Gating strategy to collect flow cytometry data**. **(A)** Bacterial populations characterized according to side scatter (SSC) and SYBR Green fluorescence (FITC). **(B)** cell counts enumerating the cells in the bacterial gate.

### Supplementary Tables

Table 1: **Culturing conditions of bacterial strains**

| Strain name | Growth media | Incubation time |
|---|---|---|
| *Pseudoalteromonas* sp. ASV16 | 10% 2216MB | 16 h[5] |
| *Pseudoalteromonas* sp. ASV39 | 10% 2216MB | 16 h[5] |
| *Alteromonas* sp. ASV76 | 10% 2216MB | 16 h[5] |
| *Alteromonas* sp. ASV109 | 10% 2216MB | 16 h[5] |
| *Vibrio anguillarum* FS144 | 100% 2216MB | 16 h + subculture for 4 h in 50% 2216MB |
| *Vibrio cyclitrophicus* ZF270 | 100% 2216MB | 16 h + subculture for 4 h in 50% 2216MB |
| *Vibrio coralliilyticus* YB1 | 100% 2216MB | 16 h + subculture for 4 h in 50% 2216MB |
| *Pseudoalteromonas* sp. AS80 | 10% 2216MB | 16 h[4] |
| *Pseudoalteromonas* sp. AS81 | 10% 2216MB | 16 h[4] |
| *Pseudoalteromonas* sp. AS85 | 10% 2216MB | 16 h[4] |
| *Pseudocolwellia* sp. G88 | 10% 2216MB | 24 h[4] |
| *Paraglaciecola* sp. AS13 | 10% 2216MB | 24 h[4] |
| *Nitrincolaceae* sp. AS21 | 10% 2216MB | 24 h[4] |
| *Shewanella* sp. AS82 | 20% 2216MB | 36 h[4] |
