## Supplementary figures and images for "Widespread attraction of marine bacteria to the macroalgae-derived polysaccharide fucoidan"

### Supplementary Video 3

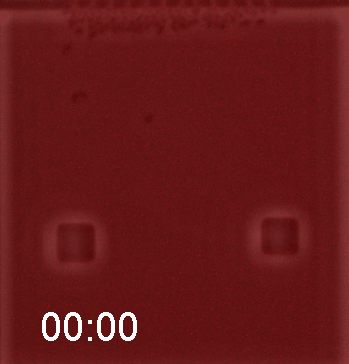
